# Glucocorticoids negatively relate to body mass on the short-term in a free-ranging ungulate

**DOI:** 10.1101/2022.09.15.508076

**Authors:** Lucas D. Lalande, Emmanuelle Gilot-Fromont, Jeffrey Carbillet, François Débias, Jeanne Duhayer, Jean-Michel Gaillard, Jean-François Lemaître, Rupert Palme, Sylvia Pardonnet, Maryline Pellerin, Benjamin Rey, Pauline Vuarin

**Affiliations:** Université de Lyon, Université Lyon 1, CNRS, Laboratoire de Biométrie et Biologie Evolutive UMR 5558, F-69622 Villeurbanne, France; Université de Lyon, VetAgro Sup, Marcy l’Etoile, France; Institute of Ecology and Earth Sciences, University of Tartu, Tartu, 51014, Estonia; Office Français de la Biodiversité, Direction de la Recherche et de l’Appui Scientifique, Service Départemental de la Haute-Garonne, Villeneuve-de-Rivière, 31800, France; Unit of Physiology, Pathophysiology and Experimental Endocrinology, Department of Biomedical Sciences, University of Veterinary Medicine, Veterinärplatz 1, 1210 Vienna, Austria; Université de Strasbourg, CNRS, IPHC UMR 7178, Strasbourg, France; Office Français de la Biodiversité, Direction de la Recherche et de l’Appui Scientifique, Service Conservation et Gestion Durable des Espèces Exploités, Châteauvillain, 52210, France

**Keywords:** stress response, body condition, Faecal Cortisol Metabolites, individual performance, ungulates

## Abstract

Environmental fluctuations force animals to adjust glucocorticoids (GCs) secretion and release to current conditions. GCs are a widely used proxy of an individual stress level. While short-term elevation in GCs is arguably beneficial for fitness components, previous studies have documented that the relationship between long-term baseline GCs elevation and fitness components can vary according to ecological and individual factors and according to the life-history of the species studied. Using longitudinal data on roe deer (*Capreolus capreolus*) from two populations facing markedly different environmental contexts, we tested whether baseline GC levels negatively correlate with body mass – a trait positively associated with demographic individual performance – on the short- to long-term. In support, higher baseline GC concentrations were associated to lighter body mass, both measured during the same capture event, in adults of both populations. Overall, we showed that despite the marked environmental and demographic differences between populations and despite the between-sex differences in life history (i.e. reproductive tactics), the relationship between body mass and GCs is consistent across environmental contexts, but might differ according to the life history stage of an individual. This work opens promising perspectives to further explore the relationship between GC and fitness-related traits according to life history stages in free-ranging mammals across seasonal and environmental contexts. The timing and context-dependence of GC levels highlight the complexity of studying stress responses in the wild.

## Introduction

Throughout their life, individuals have to adjust their morphology, physiology and/or behaviour to cope with environmental variation. Part of this variation is predictable as individuals can rely on environmental cues (*e.g.* photoperiod or temperature at the daily or seasonal scales) to anticipate future conditions (Wingfield 2003, 2008). Whereas predictable variation is not considered as stressful *per se* (*i.e.* eliciting a stress response), an unpredictable perturbation can disrupt life cycles and trigger a stress response through the activation of the hypothalamic-pituitary-adrenal (HPA) axis and the release of glucocorticoids (GCs), departing them from the baseline level (Reeder and Kramer 2005). The primary role of these hormones (namely cortisol and corticosterone) is to maintain an organism’s energy balance according to its requirements in its current environment (Wingfield 2013, Hau et al. 2016), meaning that GCs promote allostasis (*i.e.* achieving stability through change) by ensuring energy homeostasis despite predictable and unpredictable environmental and life history variation (McEwen and Wingfield 2003, 2010, Romero et al. 2009). On the short-term, an elevation of GC concentration in an individual facing a challenging situation is beneficial because it promotes processes enhancing survival through reallocation of stored energy from non-immediately essential functions (e.g. growth, reproduction) towards cardiovascular functions or locomotion and foraging activities (Sapolsky et al. 2000). After a temporary perturbation, the GC concentration often quickly returns to the baseline level (Reeder and Kramer 2005). However, baseline GC concentration may remain elevated on the long-term (*i.e.* days to weeks) as a result from exposure to repeated or long-lasting perturbations such as prolonged inclement weather or food shortage (McEwen and Wingfield 2010, Hau et al. 2016). Such elevated GC concentration in the long-term can have adverse consequences on individual performance through disruption of the immune function (Dhabhar 2014), inhibition of growth, or decreased body condition (Reeder and Kramer 2005), with an intensity depending on the species (Boonstra 2013). Indeed, long-term GC elevation can cause ‘allostatic overload’ (*i.e.* when the energy intake is lower that the energy required to support daily and seasonal activities and to cope with unpredictable perturbations, McEwen and Wingfield 2010). Although variations in GC concentration does not reflect only stress (MacDougall-Shackleton et al. 2019), GC levels are a widely used proxy for evaluating an individual physiological stress response (Palme 2019).

While many studies analysed the effect of an increase of GC beyond the baseline level (*i.e.* acute stress-induced GC concentration) on individual performance, fewer have considered how GCs at baseline concentration (*i.e.* homeostatic levels of hormones) affect the performance of animals in the wild. Results from these rare studies remain equivocal (Rogovin et al. 2003, 2008, Altmann et al. 2004, Ethan Pride 2005, Pauli and Buskirk 2007, Cabezas et al. 2007, Pedersen and Greives 2008, Pinho et al. 2019), which could be explained by the complex influence of environmental conditions (Henderson et al. 2017). When resources are limited, increased energy demands can be detrimental by resulting in high levels of GC and allostatic overload. Nevertheless, this detrimental effect may not be observed when resources are abundant, as high energy input can contribute reducing allostatic load (McEwen and Wingfield 2003, 2010, Romero et al. 2009). Food availability (*i.e.* energy availability) or behavioural flexibility stimulating foraging activity can thus compensate for elevated GC levels and associated higher energy requirements (Henderson et al. 2017). Animals inhabiting suboptimal habitats (*e.g.* exposed to human disturbance, predation, intra- or interspecific competition or pathogens) experience continuous stressful conditions and generally show a higher GC concentration compared to conspecifics inhabiting optimal areas (Dantzer et al. 2013, 2014, Formenti et al. 2018, Boudreau et al. 2019, Dulude-de Broin et al. 2020, Carbillet et al. 2020).

Additionally, there is evidence that the direction of the relationship between GCs and performance can also depend on individual attributes, such as body condition, sex or reproductive status and reproductive tactics (Tilbrook et al. 2000, Ricklefs and Wikelski 2002, Wey et al. 2015, Blumstein et al. 2016, Vuarin et al. 2019). Under high energy demands triggered by elevated GC concentration, high quality individuals are more likely to prevent themselves from reaching allostatic overload and may perform better than poor quality ones. In African striped mice (*Rhabdomys pumilio*), elevated baseline GC concentration was related to the selective disappearance of light individuals (Vuarin et al. 2019). Since the relationship between GCs and performance is complex and involves individual attributes and environmental context, both sources of variation have to be investigated to disentangle their respective roles (Crespi et al. 2013, Dantzer et al. 2014).

Focusing on body mass provides a relevant approach to investigate the detrimental effects of prolonged exposure to a high GC concentration. Indeed, body mass is often positively linked to fitness components (*e.g.* Gaillard et al., 2000; Ronget et al., 2018) and is a trait easily measured in the field that constitutes a reliable proxy of body condition in many species, especially income breeders (Hewison et al. 1996, Andersen et al. 1998). High levels of GCs are known to have a crucial role on body mass by increasing protein and amino-acid catabolism and mobilising lipids (promoting gluconeogenesis), consequently decreasing body mass (Boonstra et al. 1998, Dallman et al. 1999, Hodges et al. 2006, Rabasa and Dickson 2016).

However, the relationship between baseline GC concentration and body mass is complex. Indeed, within seasonal baseline levels, GCs actually result in increased feeding behaviours potentially promoting fat storing and body mass gain, as opposed to the previously-mentioned GC actions on protein and amino-acid catabolism and lipid mobilisation (Landys et al. 2006). Case studies have shown that the relationship between baseline GCs and body mass or condition can be positive, negative or null (*e.g.* George et al. 2014, Wey et al. 2015, Hennin et al. 2016, Boudreau et al. 2019). For instance, low body mass and poor body condition were associated with higher baseline cortisol concentrations in Eurasian badgers (*Meles meles*, George et al., 2014). On the other hand, experimental increase in GC concentration through predation risk manipulation resulted in no changes in body condition in snowshoe hares (*Lepus americanus*, Boudreau et al., 2019). Thus, more exploratory research are needed in natural settings to better understand which factors can modulate this relationship.

In this exploratory study, we aimed to contribute to a better understanding of how high baseline GC concentration relates to body mass in roe deer (*Capreolus capreolus*), an income breeder for which body mass is a good proxy for body condition (Hewison et al. 1996, Andersen et al. 2000, Pettorelli et al. 2006). We took advantage of the detailed and intensive monitoring data collected through capture-mark-recapture for more than a decade at the individual level in two populations subject to markedly different ecological contexts (*i.e.* one inhabiting a good quality habitat and one population being strongly food-limited; Pettorelli et al., 2006). To avoid measuring the capture-induced GC response (Romero & Reed, 2005), longer-term, indirect measures of GC concentration can be obtained from biological matrices other than blood (Sheriff et al. 2011). Indeed, blood GC concentration is a point in time measure and circulating GCs concentration typically increases drastically within 3 minutes following an individual’s capture (Sapolsky et al. 2000, Romero and Reed 2005), which makes it difficult to obtain reliable measures of the HPA baseline activity in free-ranging animals (but see Sheriff et al. 2011, Lavergne et al. 2021). In this context, levels of faecal glucocorticoid metabolites (FGMs) are particularly relevant as they represent an integrative measure of GC concentration several hours before the capture event (*i.e.* baseline stress; Palme, 2019). Thus, we measured FGMs, as arguably representative of baseline adrenocortical activity over a few hours, with a time delay of 12h on average in roe deer (ranging from 6 to 23 hours, Dehnhard et al. 2001).

As chronic stress can have both immediate and long-term consequences on fitness components (Monaghan and Haussmann 2015), we tested the immediate relationship between concomitant body mass and FGMs (*i.e.* short-term relationship), the relationship between FGMs in a given year and the change of body mass between two consecutive years (*i.e.* medium-term relationship) and the relationship between FGM measured on juveniles (*i.e.* individuals in their first year of life) and body mass during the prime-age and senescent life stages (*i.e.* long-term relationship). We first analysed the relationship between FGMs and body mass measured at the same capture event. We expected individuals with higher FGM concentration to be lighter than individuals with low FGM concentration. As roe deer are income breeders that do not store fat (Hewison et al. 1996, Andersen et al. 1998), we expected the catabolic actions of GCs to directly impact body mass and condition and to overcome the effects of increased feeding behaviours promoted by GCs. Then, since chronically elevated FGMs are expected to be deleterious on the medium to long-term, we tested whether FGMs measured at a given capture influenced body mass changes across lifespan. Thus, we first assessed the relationship between FGMs measured in a given year and a given body mass change between two consecutive years, separately for growing individuals and prime-aged adults having reached their full mass (*i.e.* between 4 and 10 years old; Douhard et al., 2017). We expected that higher FGM concentration should result in a smaller mass gain for growing individuals, and in a decrease in body mass for prime-aged adults, between two consecutive years, as chronic stress can be linked to inhibited growth or decreased body condition (Reeder and Kramer 2005). Finally, we tested for the effect of the stress experienced during the first year of life on the individual body mass relative to the population and sex at a given age, from the second year of life onwards. We predicted that individuals with higher FGMs during the first year of life should be lighter later in life, as late-life performance in roe deer is affected by early-life environmental conditions (Gaillard et al., 2003). Lastly, in all analyses we expected the negative relationship between FGMs and body mass to be steeper in the food-limited population due to higher allostatic load (McEwen and Wingfield 2003, Henderson et al. 2017), and we expected sexes to respond differently to higher FGMs levels due to the different reproductive tactics of males and females (Ricklefs and Wikelski 2002).

## Materials and Method

### Study population and sample collection

We studied roe deer in two populations inhabiting closed forests at Trois-Fontaines (TF – 1360 ha) in north-eastern France (48°43’N, 4°55’E) and at Chizé (CH – 2614 ha) in western France (46°05’N, 0°25’W). Roe deer are medium-sized ungulates, weighing around 25 kg and common in lowland woodlands throughout most of Europe. Roe deer display weak sexual selection, with adult males only 10% heavier than females and party size less than three females per buck. They are income breeders (Andersen et al. 2000), and to meet the markedly increased energy needs during the late gestation and early lactation periods, adult females rely on food resources rather than body reserves. Both sexes allocate high energy expenditure to reproduction. Females produce twins every year from 2 to 12 years old (Andersen et al. 1998), and males allocate heavily in territory defence for 5-6 months during the rut period (Johansson 1996). The TF population is characterised by a homogeneous habitat of good quality on a broad spatial scale, with high forest productivity due to rich soils. On the other hand, the CH population inhabits a less suitable heterogeneous habitat comprising three types of varying quality (Pettorelli et al. 2001), with low forest productivity due to poor soils and frequent summer droughts (Pettorelli et al. 2006). In both study sites, large carnivores are absent, but hunting occurs occasionally to control population growth. Since 1975 in TF and 1977 in CH, a Capture-Mark-Recapture (CMR) program has taken place every winter, and CMR analyses show that both populations were quite below carrying capacity during the present study period (2010-2021 in TF, 2013-2021 in CH, unpublished data). Captures take place during 10-12 days each year, which are spread across December (at TF) or January (at CH) and March (Gaillard et al. 1993) and consists of drive-netting captures with 30-100 beaters pushing individuals towards nets surrounding specific areas of the forests. Given that captures took place each year over 3 months, we might expect endocrine activity to vary throughout the capture season according to environmental conditions (*i.e.* populations) and sex (Dantzer et al. 2010, Sheriff et al. 2012). Therefore, we tested whether FGM levels varied as a function of the Julian date, population and sex and found that FGMs increased throughout the season in a population-specific manner (Supporting Information SI). Thus, all FGM measurements were standardised for the median Julian date of capture (9^th^ of February) specifically for each population. Successive capture events within a capture season may also have consequences on FGM measurements during the following captures. However, most capture days were more than 48 hours apart while in roe deer, FGMs peak from 6 to 23 hours after an ATCH challenge and return to baseline levels between 28 and 31 hours after treatment (Dehnhard et al. 2001). In some cases, captures took place during two consecutive days, but were then conducted on opposite areas of the forest to minimise disturbances. Likewise, morning and afternoon captures of a given day took place in different areas within both forests. We thus considered that FGM levels were not influenced by previous captures. Concerning the potential impact of the capture event itself, since roe deer were captured by drive-netting and most animal manipulations occurred between 1 and 4 hours after capture, we tested for a possible impact of this delay on FGM levels. We did not detect any increase or decrease in FGMs (log-transformed) along with the time between capture and sampling (linear regression accounting for repeated measurements of a given individual: 0.006 hour^-1^ ± 0.01 SE, t = 0.44, p = 0.7), even when the delay was > 6h (mean FGMs 6h onwards after capture compared to mean FGMs measured within 6h following capture, accounting for repeated measurement of a given individual, −0.06 ± 0.06 SE, t = −1.05, p = 0.3) (Dehnhard et al. 2001). Individuals of known age (*i.e.* captured within their first year of life) were weighed, and faecal matter has been collected since 2010. During the capture period, diet is similar and mostly composed of brambles (*Rubus* sp.) and ivy (*Hedera helix*) in the two deciduous forests (Tixier and Duncan 1996), so FGM measurements between populations should not be biased according to diet composition. During this period, females can be gestating, and differences in female reproductive status could result in FGM variability (Brunton et al. 2008, Dantzer et al. 2010). Nevertheless, we did not account for females’ reproductive status in our models, since i) we have information about reproductive status (*i.e.* pregnant or not and number of foetuses) only at CH where ultrasounds are performed on captured females, ii) most of them (*i.e.* 92 %) were gestating at that time and iii) due to the delayed embryonic implantation (Aitken 1974), gestation is at a very early stage during the capture season. Body mass measured on juveniles (*i.e.* at about 8 months of age) depends on the date of capture. Juveniles gained on average 12 g/day and 24 g/day in CH and TF, respectively, during the capture period (Douhard et al. 2017). Therefore, we standardised juvenile body mass with a linear regression, using the above-mentioned Julian date-body mass relationship. Thus, we computed the individual body mass expected on the 9^th^ of February, the median date of captures at CH and TF. In these two populations, cohort quality is reliably measured by the cohort-specific average juvenile mass corrected for the Julian date of capture (Gaillard et al. 1996). This proxy of environmental quality from the birth of an individual (typically in May) to its capture as juvenile in winter was obtained for all years, and will be referred to as ‘cohort quality’ hereafter.

### Faecal glucocorticoid metabolites (FGMs)

Baseline adrenocortical activity was estimated through FGMs (Palme 2019). We collected faecal matter rectally at capture from 2010 to 2021. Faeces were frozen at −20 °C within 24 hours prior to 2013 in CH and 2017 in TF, and immediately frozen at −80 °C after collection since then. The time between faecal sampling and freezing, as well as freezing temperature, may impact FGM values, with longer delays and higher temperature likely to result in lower FGM levels due to bacterial activity (Lexen et al. 2008, Hadinger et al. 2015, Carbillet et al. 2023b). In CH, the first FGM values available in our dataset are post-2013, so all samples were immediately frozen at −80 °C. However, in TF, FGM values are effectively higher when samples were immediately frozen (0.43 ± 0.06 SE, t = 7.62, p < 0.0001). This is accounted for in our models by adding the year of capture as a random effect. Extraction of FGM followed a methanol-based procedure and the analysis was performed using a group-specific 11-oxoaetiocholanolone enzyme immunoassay (EIA), a method previously described in details (Möstl et al. 2002) and validated for roe deer (Zbyryt et al. 2018). In brief, 0.5 g (± 0.005) of faeces were vortexed in 5 mL of 80% methanol before being centrifuged for 15 minutes at 2500 *g* (Palme et al. 2013). The amount of FGM was determined in an aliquot of the supernatant diluted 10 times in assay buffer. Measurements were done in duplicate with intra- and inter-assay coefficients lower than 10% and 15%, respectively. FGMs are expressed as nanograms per grams of wet faeces (ng/g). The data were then log-transformed for the statistical analyses (henceforth called FGMs).

### Statistical analyses

All analyses were performed using R version 4.2.2 (R Core Team 2022). Two raw FGM measures were unusually high: 5192 and 6194 ng/g, for a female in CH aged 10, and a female in TF aged 1, respectively (see SI). Results exclude these extreme values (analyses including them are reported in SI), but we specify whether adding them yielded different conclusions or not.

#### FGM repeatability

Individual FGM repeatability was calculated for all individuals, and separately for each population, using the ‘rptR’ package (Stoffel et al. 2017). Repeatability analysis included individuals sampled only once to improve estimates of the within-individual variance (Martin et al. 2011). Within-individual FGM repeatability was detectable but weak in both CH (r = 0.14, 95% CI = [0.03, 0.26]) and TF (r = 0.15 [0.01, 0.29]).

#### Model structure

We tested the hypothesis that body mass is related to FGM levels, and that this relationship may depend on individual (*i.e.* sex) and environmental (*i.e.* population and condition at birth) characteristics, using linear mixed effect models (LMMs) with a normal error distribution. All continuous covariates were mean-centred so that the intercept is the relative individual body mass for the mean cohort quality and/or mean FGMs. Visual inspection of model residuals confirmed that they fulfilled the assumptions of Gaussian distribution and homoscedasticity.

#### Short-term relationships between FGMs and body mass measured at the same capture event

We first assessed the relationship between FGMs and body mass measured the same year (*i.e.* at the same capture) to test the hypothesis that higher baseline FGMs result in short-term adverse consequences on body mass. Individuals in their first year of life (*i.e* juveniles) were analysed separately from individuals in their second year of life onwards, because juveniles have to allocate to growth and are much more susceptible to any environmental harshness than adults, making the first year of life is the critical period of roe deer population dynamics (Hamel et al. 2009, Gaillard et al. 2013). In addition, metabolic rate is generally higher in juveniles than in adults due to the costs of growth (Glazier 2005) and metabolic rates can alter hormones metabolization and excretion, making it difficult to compare juvenile and adult FGM concentrations (Goymann 2012). Since juveniles are not yet reproductively active, they also likely display different endocrine profiles than adults (Dantzer et al. 2010). Finally, adult roe deer habituate faster than juveniles to stress, which can in turn affect the relationship between FGM and fitness-related trait (Bonnot et al. 2018). Body mass varies with age, and at a given age, varies between sexes and according to environmental conditions at birth (*i.e.* cohort effects, Hamel et al. 2016). We thus tested the relationship between FGMs measured in a given year and individual body mass relative to the average body mass of all individuals of the same age, population and sex, considering only ages for which we had data from at least 3 individuals. Therefore, the average body mass was calculated up to 10 and 15 years old in males and females at CH, respectively, and up to 12 and 13 years old in males and females at TF, respectively. For juveniles, we analysed the relationship between FGMs and body mass measured the same year on 368 juveniles (78 females and 96 males in CH, 91 females and 103 males in TF). The response variable was the mass corrected for the date of capture (see above) and fixed explanatory variables included FGMs (corrected for the date of capture and the population), cohort quality, population, sex and the two-way interactions between FGMs and the other covariables. Cohort quality was expressed as a relative cohort quality (*i.e.* the difference between mean cohort quality in each population and an individual cohort quality), so there is no redundancy with the population variable. The year of birth of the individuals (*i.e.* cohort) was included as a random effect. For individuals aged 2 years or older, the dataset included 655 observations on 377 individuals: 104 females (218 observations) and 83 males (136 obs.) in CH, and 106 females (164 obs.) and 84 males (137 obs.) in TF. The response variable used to test for the FGMs-mass relationship was the individual body mass minus the average mass of roe deer of the same population, same sex and same age, regardless of their year of capture (hereafter ‘relative body mass). Fixed effects included continuous variation in FGMs, relative cohort quality, population, sex and the two-way interactions between FGMs and other covariables. Random effects of the cohort and of the year of capture were also included. Random effect of the individual identity (ID) was also included to account for repeated measurements on the same individuals.

#### Medium-term relationships between FGMs and body mass change between two consecutive years

Baseline FGMs can reflect medium-term consequences of GCs on body mass. The relationship between FGMs and the change of body mass between two consecutive years was analysed separately for early-growing individuals (*i.e.* between age 1 and 2), late-growing individuals (*i.e.* between the second year of life and adulthood at 4 years old, Hewison et al. 2011) and for prime-aged adults which had reached their full body mass (*i.e.* from 4 to 10 years old, Hewison et al., 2011). The response variable was the change in relative body mass. Briefly, the change in relative mass expresses the change in body mass of a given individual between two ages in relation to the change in body mass of all individuals of the same sex and population between the same ages. The relative mass was calculated as the difference between the mass of an individual at age *x*, and the mean mass of all individuals of age *t* of the same sex and the same population. The change in relative body mass was calculated as the difference between the relative mass at age *t* + 1 and the relative mass at age *t*. Fixed effects included relative cohort quality, population, sex, FGMs (either measured at age *t* or considered as the average of the FGM values measured at age *t* and age *t* + 1), body mass at age *t* (*i.e.* initial body mass) and all two-way interactions between FGMs and the other covariables. Random effect of the cohort was included, and for late-growing and prime-aged individuals, random effects of the year of capture and individual ID were also included. For growing individuals, the dataset included 99 individuals with FGMs measured as juveniles, and 67 individuals when including FGMs measured during the first and second years of life to calculate the mean value. For both late-growing individuals and adults, we kept a single observation per individual since the inclusion of individual ID as a random effect created singularities. For individuals with several observations, we kept the one for which the individual’s age was the closest to the mean age of all individuals with a unique observation according to sex and population, among observations with complete data. The final dataset comprised 85 individuals in their late-growth period and 90 prime-aged adults with FGMs measured at age *t*, and 63 late-growing individuals and 71 prime-aged adults for which we had measurements at age *t* and *t+1*.

#### Long-term relationships between FGMs during early life and body mass later in life

As elevated baseline FGMs are expected to have negative effects on body mass on the long-term, and as environmental conditions during the first year of life have carry-over effects on performance later in life in roe deer (Gaillard et al. 2003), we tested whether FGMs during development are associated with adult body mass. Only individuals for which we measured FGMs as juvenile and body mass beyond the second year of life were analysed. We used 345 observations on 159 individuals for which we measured FGMs as juvenile and body mass as adult: 34 females (78 obs.) and 44 males (87 obs.) in CH, and 42 females (99 obs.) and 39 males (81 obs.) in TF. The response variable was the relative individual body mass as defined above. Fixed effects included FGMs measured when juvenile, relative cohort quality, population, sex and the two-way interactions between FGMs and the other covariables. Random effects of the cohort and the year of capture were included, as well as individual ID to account for repeated measurements on the same individuals.

#### Model selection

Final models were selected based on the second order Akaike Information Criterion (AICc, *i.e.* Akaike Information Criterion corrected for small sample sizes). We compared all sub-models included in the full model described above and for each model AICc scores were computed with the “*MuMIn*” package (v. 1.47.5, Bartoń 2023). We retained the best-fitting model as the one with the lowest AICc score (Burnham and Anderson 2002), or the simplest model (*i.e.* with the lowest number of parameters) within the set of models within 7 ΔAICc. Indeed, models within 7 ΔAICc are plausible models, become increasingly equivocal up to 14 ΔAICc, and implausible afterwards (Burnham et al. 2011). For each variable we estimated its effect size (β) with 95% confidence intervals (95% CI) and calculated marginal and conditional R². We provide full model selection tables in SI.

## Results

FGM concentrations ranged from 8 to 3428 ng/g with a median value of 689 ng/g. The median FGMs was 674 ng/g (range: 34-3275 ng/g) in CH, and 715 ng/g (range: 8-3428 ng/g) in TF.

### Short-term relationships between FGMs and body mass measured at the same capture event

#### Juveniles

The retained model highlighted the expected positive relationship between cohort quality and body mass (β_Cohort quality_ = 1.08 [0.84, 1.32], Table 1, SI), but no relationship between FGMs and body mass could be evidenced (Figure 1(a), Table 1, SI). Results were similar when including female of TF with a high FGM value (SI). Note that since the random effect of the cohort created singularities, it was removed and models therefore consisted of linear models.

**Figure 1.**
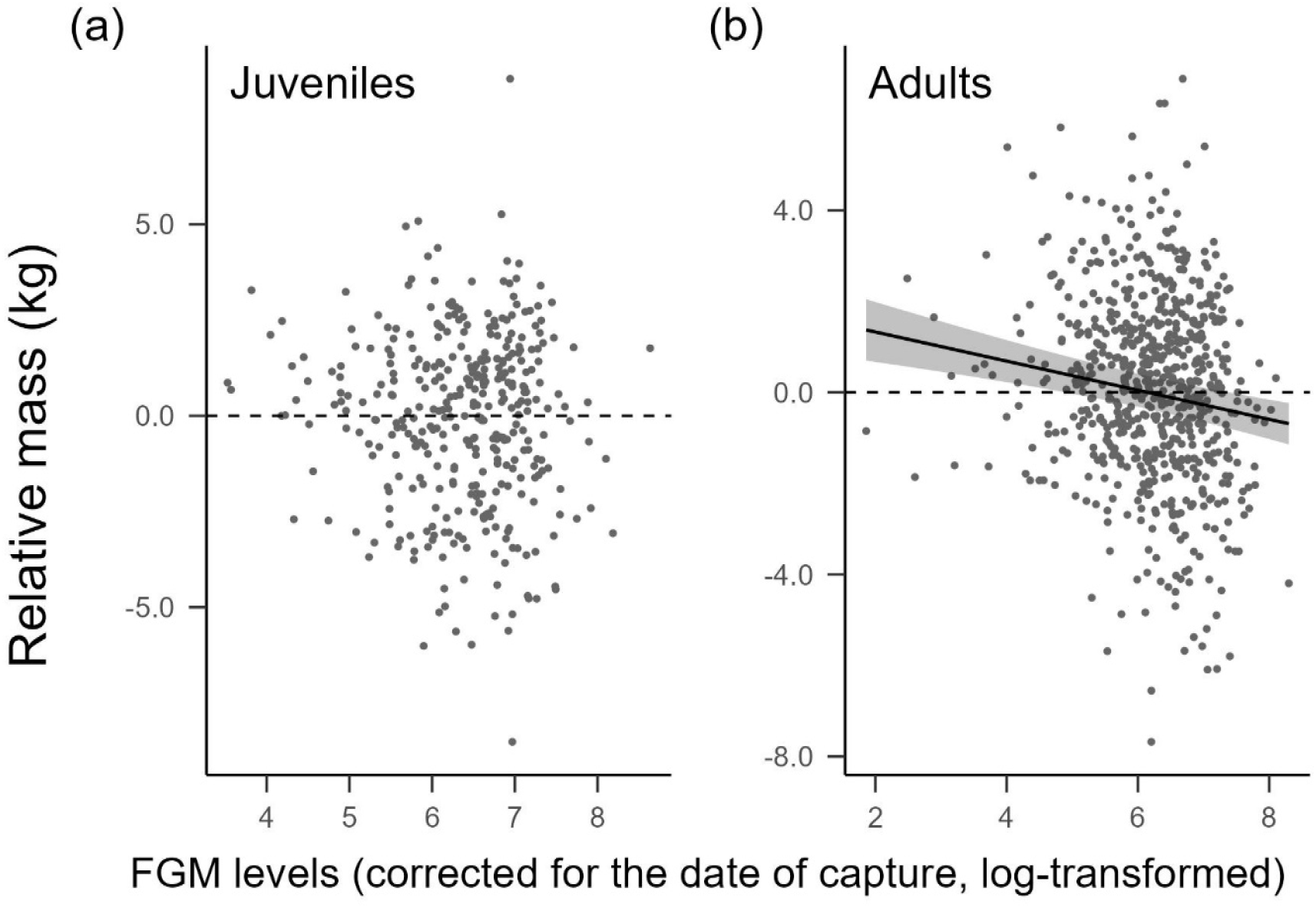
Relationship between faecal glucocorticoid metabolites (log-transformed FGMs, corrected for the date of capture and population) and individual relative body mass (individual body mass (corrected for the date of capture for juveniles) minus the average body mass of all individuals of the same age, population and sex) for (a) juveniles (N = 368 individuals) and for (b) adults aged 2 years old onwards (N = 655 observations on 377 individuals). Points are raw data and line is the prediction from the retained model with 95% confidence intervals.

**Table 1.** Linear and linear mixed effect models selected for the short-, medium- and long-term relationships between relative body mass and faecal glucocorticoid metabolites (FGMs). Models accounted for sex, population and for relative cohort quality (Qcohort, the difference between the mean cohort quality in each population and an individual cohort quality). Models were selected through model selection based on AICc. 95%CI: 95% confidence intervals, V: variance, SD: standard-deviation.

| SHORT-TERM |  |  |  |  |  |  |  |  |
| --- | --- | --- | --- | --- | --- | --- | --- | --- |
| Juveniles |  |  | Adults |  |  |  |  |  |
| Random effects | V | SD | Random effects | V | SD |  |  |  |
|  |  |  | Individual ID | 3.45 | 1.86 |  |  |  |
|  |  |  | Year of capture | 0.29 | 0.54 |  |  |  |
| Fixed effects | Estimate | 95%CI | Fixed effects | Estimate | 95%CI |  |  |  |
| Intercept | -0.06 | [-0.28, 0.16] | Intercept | -0.02 | [-0.39, 0.35] |  |  |  |
| Qcohort | 1.08 | [0.84, 1.32] | FGM | -0.32 | [-0.46, -0.18] |  |  |  |
| Marginal R<sup>2</sup> | 0.18 |  | Marginal R<sup>2</sup> | 0.02 |  |  |  |  |
| Conditional R<sup>2</sup> |  |  | Conditional R<sup>2</sup> | 0.81 |  |  |  |  |
| MEDIUM-TERM |  |  |  |  |  |  |  |  |
| FGM measured at age *t* |  |  |  |  |  |  |  |  |
| Early-growth |  |  | Late-growth |  |  | Prime-aged |  |  |
| Random effects | V | SD | Random effects | V | SD | Random effects | V | SD |
| Cohort | 0.65 | 0.81 | Year of capture | 0.41 | 0.64 | Year of capture | 0.24 | 0.49 |
| Fixed effects | Estimate | 95%CI | Fixed effects | Estimate | 95%CI | Fixed effects | Estimate | 95%CI |
| Intercept | -0.35 | [-0.92, 0.22] | Intercept | -0.55 | [-1.02, -0.08] | Intercept | -0.19 | [-0.60, 0.22] |
|  |  |  | Initial mass | -0.15 | [-0.25, -0.05] |  |  |  |
| Marginal R<sup>2</sup> | - |  | Marginal R<sup>2</sup> | 0.09 |  | Marginal R<sup>2</sup> | - |  |
| Conditional R<sup>2</sup> | 0.21 |  | Conditional R<sup>2</sup> | 0.32 |  | Conditional R<sup>2</sup> | 0.15 |  |
| Mean FGM (FGM<sub>*t*</sub> and FGM<sub>*t+1*</sub>) |  |  |  |  |  |  |  |  |
| Early-growth |  |  | Late-growth |  |  | Prime-aged |  |  |
| Random effects | V | SD | Random effects | V | SD | Random effects | V | SD |
| Cohort | 0.90 | 0.95 | Year of capture | 0.32 | 0.56 | Year of capture | 0.18 | 0.42 |
| Fixed effects | Estimate | 95%CI | Fixed effects | Estimate | 95%CI | Fixed effects | Estimate | 95%CI |
| Intercept | -0.37 | [-1.11, 0.37] | Intercept | -0.47 | [-0.94, -0.00] | Intercept | -0.28 | [-0.71, 0.15] |
|  |  |  | Initial mass | -0.17 | [-0.27, -0.07] |  |  |  |
| Marginal R<sup>2</sup> | - |  | Marginal R<sup>2</sup> | 0.13 |  | Marginal R<sup>2</sup> | - |  |
| Conditional R<sup>2</sup> | 0.26 |  | Conditional R<sup>2</sup> | 0.32 |  | Conditional R<sup>2</sup> | 0.10 |  |
| LONG-TERM |  |  |  |  |  |  |  |  |
| Random effects |  |  | V |  |  | SD |  |  |
| Individual ID |  |  | 3.65 |  |  | 1.91 |  |  |
| Year of capture |  |  | 0.63 |  |  | 0.79 |  |  |

| Fixed effects | Estimate | 95%CI |
| --- | --- | --- |
| Intercept | -0.06 | [-0.63, 0.51] |
| Marginal R<sup>2</sup> | - |  |
| Conditional R<sup>2</sup> | 0.83 |  |

#### Adults

We found support for a short-term negative relationship between individual relative body mass and FGMs in adults (β_FGM_ = −0.32 [-0.46, −0.18], Figure 1(b), Table 1, SI). Results were similar when including the high FGM value measured on a female of CH (SI). Since the random effect of the cohort created singularities, the models only included individual ID and year of capture as random effects.

### Medium-term relationships between FGMs and change in body mass between two consecutive years

#### Early-growing individuals

We did not find any evidence for a relationship between FGMs, either measured in juveniles or considered as the mean FGM value between the first and second years of life, and the change in relative body mass for growing individuals (Figure 2(a), Table 1, SI). Results were similar when including the female of CH with the highest FGM value (SI). All models included cohort as random effect.

**Figure 2.**
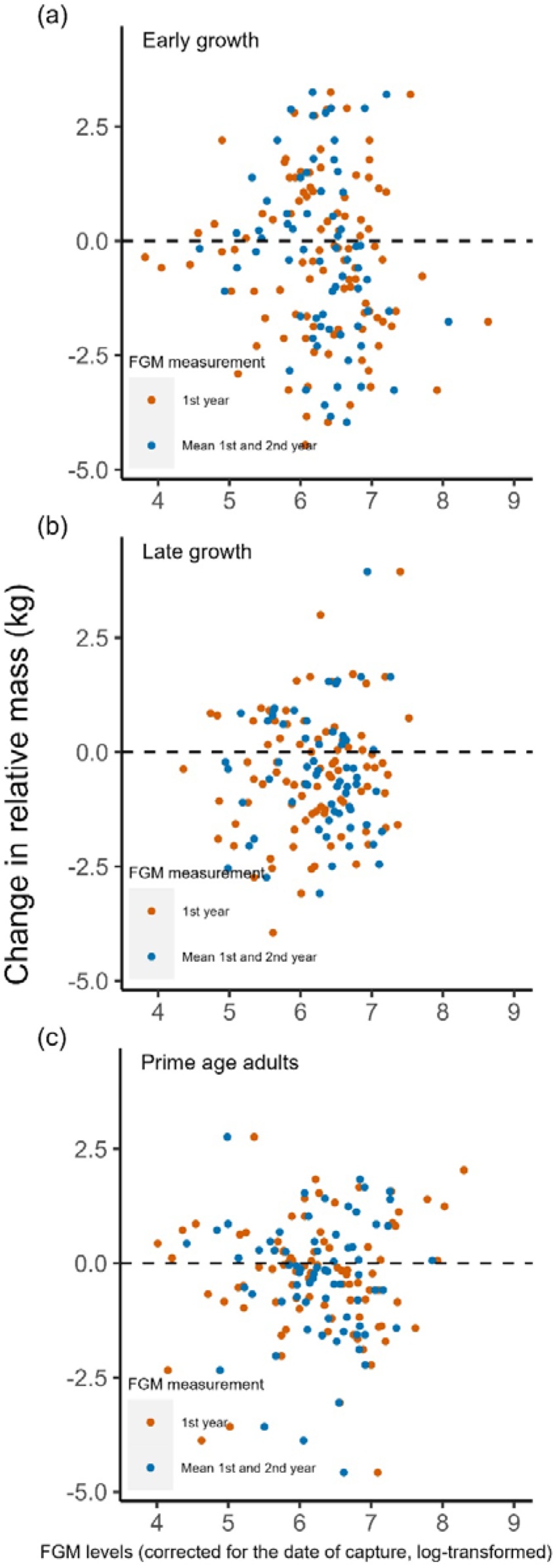
Relationship between faecal glucocorticoid metabolites (log-transformed FGMs, corrected for the date of capture and population) and change in relative body mass between two consecutive years for (a) early-growing individuals (between the first and second years of life), with FGMs either measured the first year of life (N = 99, orange points) or considered as the mean FGMs between the first and second years (N = 67, blue points); for (b) late-growing individuals (between 2 and 4 years old) with FGMs either measured at age *t* (N = 85, orange points) or considered as the mean FGMs between years *t* and *t+1* (N = 63, blue points) and for (c) prime-aged adults which had reached their full body mass (from 4 years old to 10 years old), with FGMs either measured at a given age *t* (N = 90, orange points) or considered as the mean FGM between age *t* and age *t+1* (N = 71, blue points). Points are raw data

#### Late-growing individuals

We found a negative relationship between body mass gain and initial mass (β_mass_ = −0.15 [-0.25, −0.05], Table 1, SI) for the dataset accounting for FGMs measured in juveniles. With the dataset including the mean FGM value between FGMs measured at age *t* and age *t+1*, we also found a negative relationship between late growth and body mass measured at age *t* (β_mass_ = −0.17 [-0.27, −0.07], Table 1, SI). However, in both cases, we found no support for a relationship between FGMs and change in relative body mass (Figure 2(b), Table 1, SI). Models resulted in singularities when the random effect of the cohort was included so that only the year of capture was included as a random effect in the models.

#### Prime-aged adults

In no cases (*i.e.* neither when accounting for FGM measured at age *t* or for the mean FGM value between age *t* and age *t+1*) was a relationship found between FGMs and change in relative body mass (Figure 2(c), Table 1, SI). Similarly, we found no evidence that mass measured at age *t*, cohort quality, population or sex were related to changes in relative body mass (Table 1, SI). Models resulted in singularities when the random effect of the cohort was included, thus, only the random effect of the year of capture was included in the models.

### Long-term relationships between FGMs during early-life and body mass later in life

We found no evidence for a relationship between FGM measured in juveniles, cohort quality, population or sex, and adult relative body mass (Figure 3, Table 1, SI). Results were similar when including the three observations of the adult female of CH with the highest FGM value measured as juvenile (SI). We only included individual ID and year of capture as random effects, because including the cohort created singularities.

**Figure 3.**
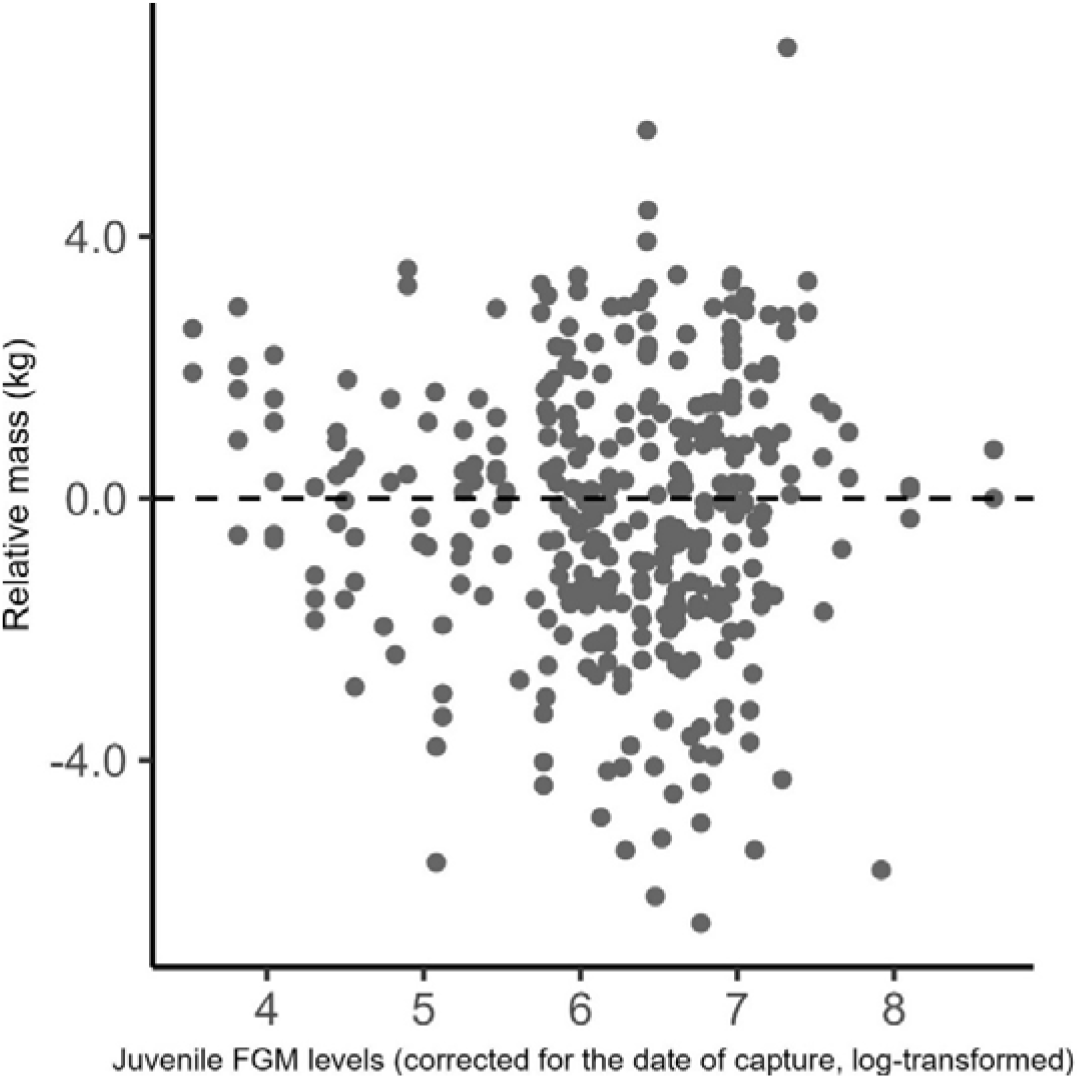
Relationship between faecal glucocorticoid metabolites (log-transformed FGMs, corrected for the date of capture and population) measured in juveniles and individual relative body mass (*i.e.* individual body mass minus the average body mass of all individuals of the same age, sex and population) (N = 345 observations on 159 individuals). Points are raw data.

## Discussion

The relationship between GCs and fitness-related traits such as body mass is yet to be fully deciphered. Recent reviews indicate that studies in the wild have led to contrasting results by showing either negative, positive or no relationships between measures of stress and studied traits (Bonier et al. 2009, Crespi et al. 2013, Dantzer et al. 2014). A salient conclusion from these studies is that the relationship depends on an individual’s ecological context and internal state. In the present study, the longitudinal monitoring of individual roe deer in two populations exposed to markedly different environmental contexts – one facing limiting resources and the other benefiting from a higher availability and quality of resources – allowed us to analyse the relationship between FGMs and body mass, while accounting for ecological factors and individual attributes such as sex and age. On the short-term, we found clear evidence for a negative relationship between baseline adult FGMs and adult body mass, an important driver of fitness in this species (Gaillard et al. 2000).

We could not detect any relationship between FGMs and the change in relative body mass between two consecutive years, neither in growing or prime-aged individuals. In late-growing individuals only, a negative relationship between initial body mass and relative body mass gain appeared, so that lighter individuals of age *t* tended to gain more weight from one year to the other, suggesting the existence of compensatory growth mechanisms in these populations (Metcalfe and Monaghan 2001). Likewise, we found no support for a relationship between early-life FGM levels and adult body mass. Indeed, we found that FGMs and body mass were negatively associated only on the short-term and in adult individuals only. This suggest that FGMs relate to body mass, but without any carry-over effects, and that FGMs are related to body mass differently according to life-history stages. High viability selection at the juvenile stage in the two populations (Toïgo et al. 2006, Douhard et al. 2014) could explain that we could not observe any carry-over effects of FGMs on body mass, and an absence of evidence for a short-term relationship between FGMs and body mass in juveniles.

Roe deer body mass correlated negatively with FGM levels on the short-term, in adults, with no detectable population or sex differences. This result suggests that the relationship between FGM and body mass is consistent across individuals and between environmental contexts in this species. Roe deer are income breeders, relying mostly on acquired resources rather than on stored energy (Andersen et al. 2000). As GCs stimulate activity through the mobilisation of energy stored (Sapolsky et al. 2000, Reeder and Kramer 2005), the physiological costs of elevated GC levels are expected to directly impair body condition as individuals do not have body reserves to buffer elevated energy requirements (Henderson et al. 2017). However, given marked between-population differences in environmental context we would have expected individuals at TF to be able to mitigate the physiological costs of high GC levels by reducing their allostatic load (McEwen and Wingfield 2010, Henderson et al. 2017), compared to individuals at CH which were expected to enter allostatic overload, thus expecting different allostatic load in the two populations (McEwen and Wingfield 2010). Both populations differ in terms of overall habitat quality, with CH being characterised by a low forest productivity due to poor soils, mild winters, but frequent droughts during summer, and TF by a high forest productivity due to rich soils and relatively wet summers (Pettorelli et al. 2006). Accordingly, life history parameters differ between populations. This is especially the case for juvenile survival (Gaillard et al. 1993), body condition (Gilot-Fromont et al. 2012), senescence patterns (haematological parameters: Jégo et al. 2014, body mass: Douhard et al. 2017, immune system: Cheynel et al. 2017), and the relation between FGMs and immune traits (Carbillet et al. 2023a). Thus, differences in the relationship between FGMs and body mass was expected to occur between populations, with the strongest negative association in CH, the poorer habitat in which individuals would be more at risk of allostatic overload (McEwen and Wingfield 2010, Henderson et al. 2017). Red deer (*Cervus elaphus*) showed different allostatic loads across populations with different densities and environmental conditions, with individuals facing high density or poor-quality habitat having higher FGM levels and lower body mass (Caslini et al. 2016). Our results rather suggest that both roe deer populations are in a state of allostatic overload. During the winter season, roe deer face more severe winters at TF than at CH, which can in turn mitigate, during this period, the overall better resource quality in TF. This could partly explain why no population differences were detected. It also suggests that allostatic load can change across seasons, as shown in wild grey mouse lemurs (*Microcebus murinus*, Hämäläinen et al. 2015), and that this variation could operate in a population-specific way. It is therefore of great interest to assess the GC-fitness relationship across different environmental contexts, including different years or climatic conditions, and through various life history stages. Finally, the absence of large predators and the weak hunting pressure in both population might at least partly account for the absence of support for a steeper relationship at CH. Studies of roe deer populations facing varying conditions in terms of presence/absence of large predators and hunting pressure, in Poland, showed that hunted populations without large predators had highest FGM levels with largest variation (Zbyryt et al. 2018). In support, observed variation in FGM levels was high in both populations (CV = 59.7 % and 55.3 % in CH and TF, respectively), which could mitigate the relationship between body mass and FGMs and differences between populations.

Similarly, we could not detect any sex difference, despite the marked differences in the stress response in males and females reported in mammals, due in part to the interaction between the HPA and reproductive axes (Viau 2002, Wilson et al. 2005, and see Toufexis et al. 2014, Novais et al. 2016 for reviews). This absence of sex-differences also suggests that during this period of the year, female reproductive status does not influence the relationship between FGMs and body mass. In reindeer (*Rangifer tarandus*), cortisol levels did not differ between males, non-pregnant or pregnant females, but showed seasonal variation (Bubenik et al. 1998). Accordingly, red deer showed no sex-differences in their levels of FGMs but also showed seasonal variation (Huber et al. 2003). Thus, it would be interesting to collect FGM data of roe deer during different seasons as the stress response can vary accordingly (Bubenik et al. 1998, Vera et al. 2011). It would also provide an opportunity to investigate whether the different reproductive tactics of males and females can affect the relationship between GC and fitness-related traits. Females of an iteroparous, long-lived species such as roe deer are expected to trade reproduction for survival when environmental conditions are poor (Hirshfield and Tinkle 1975). Thus, females are expected to be stress-responsive during their reproductive period to be able to switch between alternative physiological states according to whether they reproduce or not, due to their future opportunities of reproducing (Ricklefs and Wikelski 2002). On the other hand, males allocate heavily to reproduction by defending territories for half of the year (Johansson 1996). Territory defence results in elevated physiological damages, whatever their reproductive success. Therefore, females could show a larger variability in the way they respond to GC than males which are more constrained. This provides promising perspectives to better understand the relationship between GC levels and fitness-related traits according to sex-specific life history stages. It would also be interesting to investigate whether offspring have different life-history trajectories according to their mothers’ baseline FGMs during gestation, as shown in North American red squirrels (*Tamiasciurus hudsonicus*, Dantzer et al. 2013), or according to whether mother were captured or not during gestation to further investigate the impact of capture on individuals.

FGMs have been previously used as proxies to evaluate the homeostatic level or allostatic load in wild populations, but the results should be interpreted with caution. FGMs do not fully reflect an individual’s stress response because GCs only represent one part of the complex endocrine pathway involved in this response (MacDougall-Shackleton et al. 2019). Moreover, although GCs are often referred to as “stress hormones”, at baseline levels, their primary role is to acquire, deposit and mobilise energy (Busch and Hayward 2009). Thus, unlike acute GC concentrations, baseline GCs do not only provide information on how an individual responds to stressors, but also on its physiology and activity (Reeder and Kramer 2005). It has been reported that the acute GC response tends to be attenuated in individuals repeatedly exposed to stressors (*e.g.* Ader et al. 1968, Kant et al. 1983, Vera et al. 2011). However, baseline GC levels can also, but not always, be lower in individuals chronically exposed to environmental stressors, making it difficult to interpret FGM levels in the wild. Indeed, whether low baseline GCs actually represent a non-stressed individual rather than an individual chronically exposed to stressors is difficult to tell (Davis and Maney 2018).

In conclusion, we found that high baseline FGMs may have immediate adverse consequences on body mass and that the relationship between body mass and FGM levels seems to depend on an individual life-stage, rather than on the environmental context or sex. Through this work, we therefore emphasise the need to account for both environmental and individual factors, including life history traits, to better capture the relationship between the stress response and fitness-related traits. We also put emphasis on the idea to collect FGM across different temporal and environmental contexts to evaluate how the GC-fitness relationship can be modulated according to the seasonal and environmental conditions. To do so, it appears critical to have access to longitudinal data. Measuring proxies of stress together with fitness-related traits throughout the life of individuals is likely to be the only way to properly refine our understanding of the implications of stress for fitness.

## Ethics

The research presented in this manuscript was done according to all institutional and/or national guidelines. For both populations (Trois-Fontaines and Chizé), the protocol of capture of roe deer, under the authority of the Office Français de la Biodiversité (OFB), was approved by the Director of Food, Agriculture and Forest (Prefectoral order 2009-14 from Paris). All procedures were approved by the Ethical Committee of Lyon 1 University (project DR2014-09, June 5, 2014).

## Acknowledgements

This work was conducted as part of a Ph.D. funded by the Université de Lyon and the Office Français de la Biodiversité (OFB). We are grateful to all technicians, researchers and volunteers participating in the collect of data on both sites (Trois-Fontaines and Chizé). We particularly thank the editor and two anonymous reviewers for their insights and valuable comments, and Philippe Veber and Marie-Laure Delignette-Muller for their precious help considering the statistical analyses.

