## Supporting Information for "Glucocorticoids negatively relate to body mass on the short-term in a free-ranging ungulate"

[6] Current institution: Université de Strasbourg, CNRS, IPHC UMR 7178, Strasbourg, France

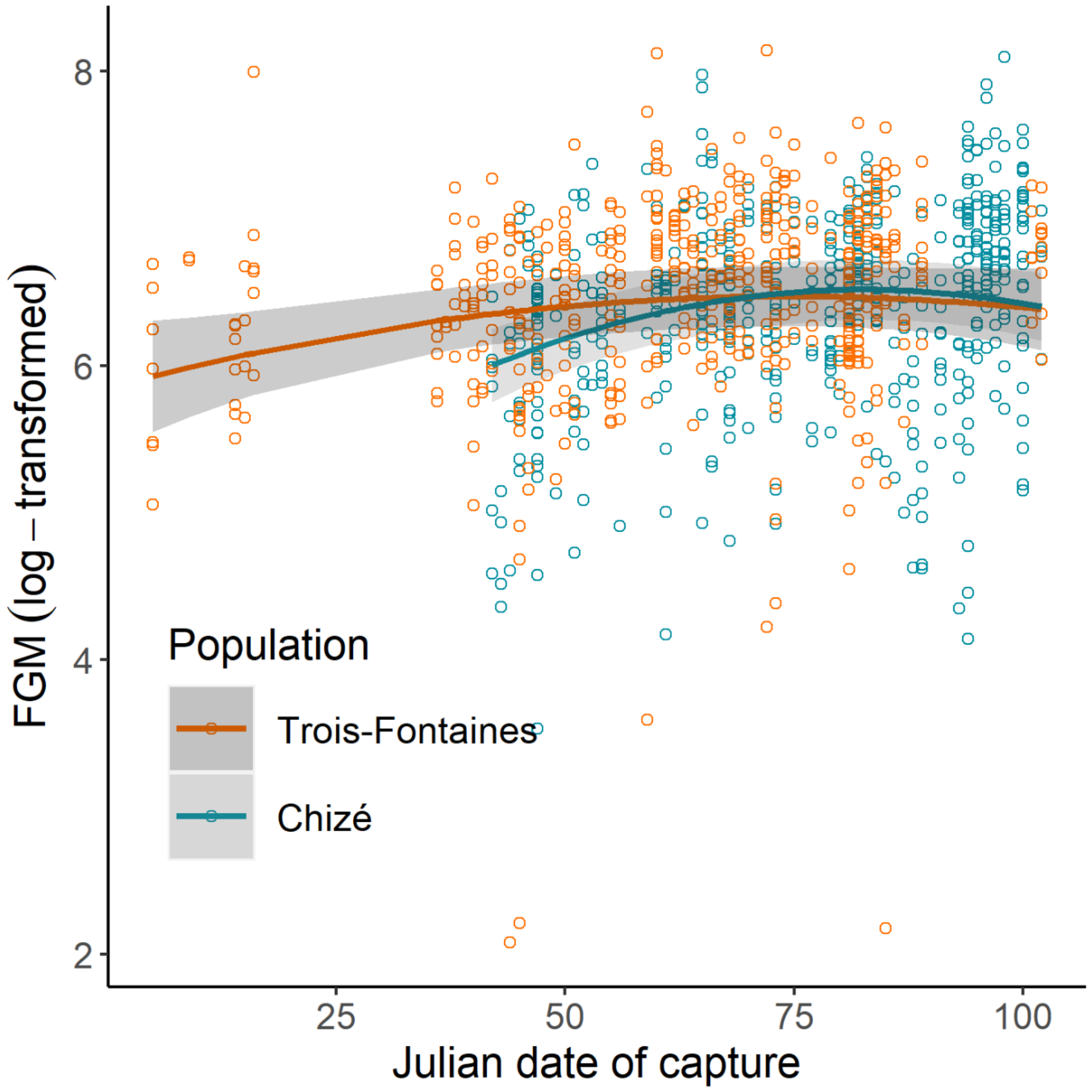


**Figure S1.** Seasonal variation of faecal glucocorticoid metabolites (FGMs, log-transformed) according to Julian date of capture (starting early December, ending early March) and the population (Trois-Fontaines: orange, Chizé: blue). FGMs increase across the capture season and slightly decrease at the end of the season, in a population-specific manner. Points are raw data and lines are predictions from the selected model with 95% confidence intervals.

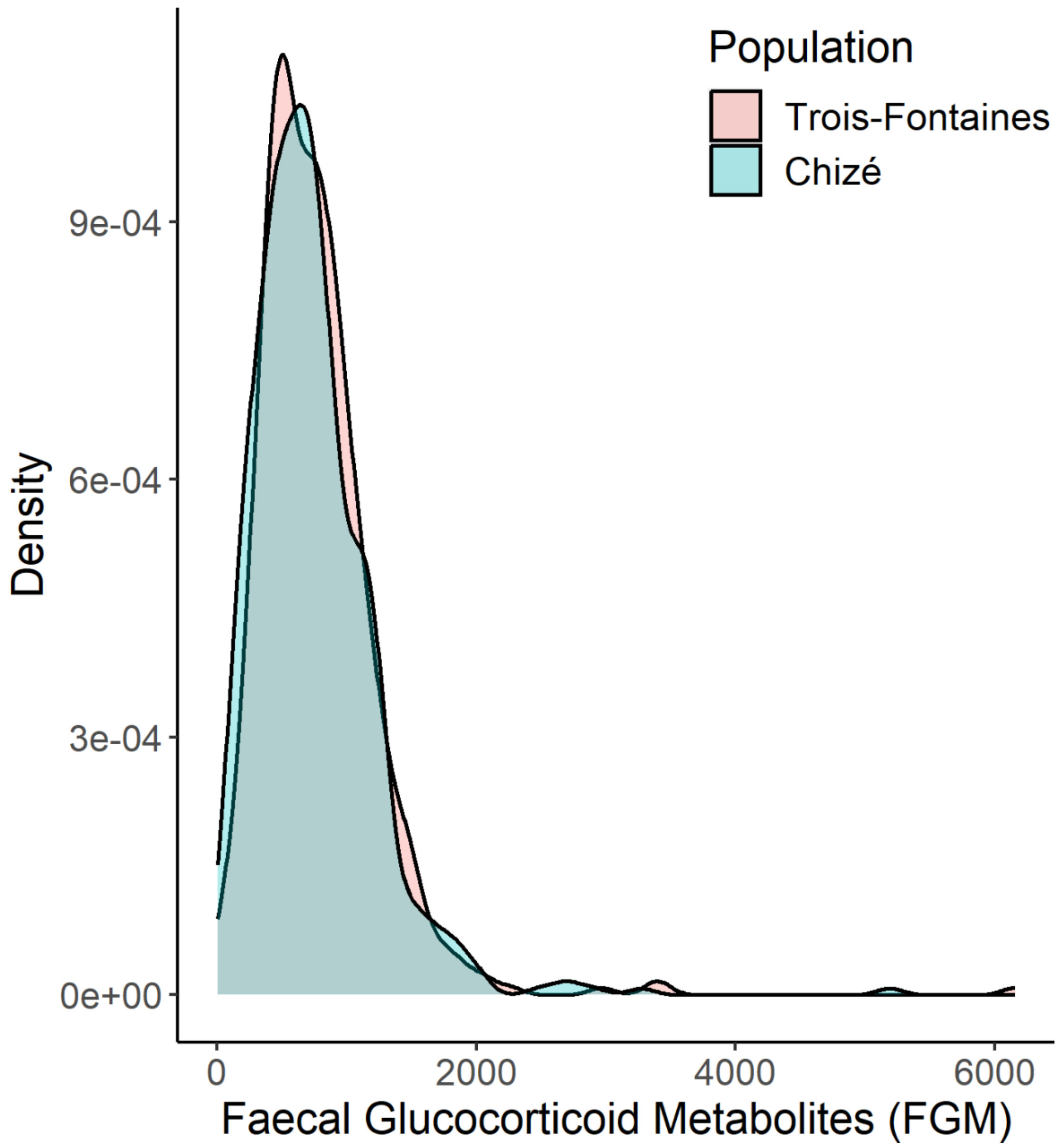


**Figure S2.** Density plot of faecal glucocorticoid metabolites (FGMs) according to population, evidencing particularly high FGM values.

**Table S1.** Linear and linear mixed effect models selected for the short-, medium- and long-term relationships between relative body mass and faecal glucocorticoid metabolites (FGM), including two high FGM values. Models accounted for sex, population (CH: Chizé) and for cohort quality (Qcoh). Models were selected through model selection based on AICc and parameters estimated through model averaging. 95%CI: 95% confidence interval, V: variance, SD: standard-deviation.

| SHORT-TERM | | | | | |
|---|---|---|---|---|---|
| Juveniles | | | Adults | | |
| **Random effects** | **V** | **SD** | **Random effects** | **V** | **SD** |
| | | | Individual ID | 3.45 | 1.86 |
| | | | Year of capture | 0.29 | 0.53 |
| **Fixed effects** | **Estimate** | **95% CI** | **Fixed effects** | **Estimate** | **95% CI** |
| Intercept | -0.06 | [-0.28, 0.16] | Intercept | -0.02 | [-0.39, 0.35] |
| Qcoh | 1.08 | [0.84, 1.32] | FGM | -0.33 | [-0.47, -0.19] |
| Marginal $R^2$ | 0.18 | | Marginal $R^2$ | 0.02 | |
| Conditional $R^2$ | | | Conditional $R^2$ | 0.81 | |

| MEDIUM-TERM (Early-growth) | | | | | |
|---|---|---|---|---|---|
| FGM measured year *t* | | | Mean FGM (FGM*t* and FGM*t+1*) | | |
| **Random effects** | **V** | **SD** | **Random effects** | **V** | **SD** |
| Cohort | 0.72 | 0.85 | Cohort | 0.99 | 0.99 |
| **Fixed effects** | **Estimate** | **95%CI** | **Fixed effects** | **Estimate** | **95% CI** |
| Intercept | -0.32 | [-0.91, 0.27] | Intercept | -0.33 | [-1.09, 0.43] |
| Marginal $R^2$ | - | | Marginal $R^2$ | - | |
| Conditional $R^2$ | 0.22 | | Conditional $R^2$ | 0.27 | |

| LONG-TERM | | |
|---|---|---|
| **Random effects** | **V** | **Random effects** |
| Individual ID | 3.69 | 1.92 |
| Year of capture | 0.65 | 0.80 |
| **Fixed effects** | **Estimate** | **95% CI** |
| Intercept | -0.04 | [-0.61, 0.53] |
| Marginal $R^2$ | - | |
| Conditional $R^2$ | 0.83 | |

Table S2. Linear mixed effects model (LMM) selection table for the short-term relationship between relative body mass for juveniles and adults (*i.e.* aged 2 years old and older) and concomitant faecal glucocorticoid metabolites (‘FGM’). We accounted for cohort quality (‘Qcoh’), population (reference level is Chizé: CH, and is compared to Trois-Fontaines: TF), and sex (reference level is male: M, and is compared to females: F). Values give the parameter coefficient and values between brackets in grey rows are the standard-errors. ‘I’ is the Intercept, ‘df’ is the number of parameters, ‘Log-lik’ is the log-likelihood, ‘delta’ is the difference of AICc between the candidate model and the model having the lowest AICc, and ‘weight’ the AIC weight of each model. Retained model is in bold. Only models up to delta AICc = 7 are displayed.

| SHORT-TERM (JUVENILES, N = 368) | | | | | | | | | | | | |
|---|---|---|---|---|---|---|---|---|---|---|---|---|
| I | FGM | Qcoh | Pop(CH) | Sex(M) | FGM:Qcoh | FGM:Pop(CH) | FGM:Sex(M) | df | Log-lik | AICc | Delta | Weight |
| -0.06 | -0.23 | 1.09 | | | | | | 4 | -785.24 | 1578.59 | 0.00 | 0.23 |
| (0.11) | (0.13) | (0.12) | | | | | | | | | | |
| **-0.06** | | **1.08** | | | | | | **3** | **-786.72** | **1579.52** | **0.93** | **0.15** |
| **(0.11)** | | **(0.12)** | | | | | | | | | | |
| -0.06 | -0.22 | 1.08 | | | -0.07 | | | 5 | -785.16 | 1580.49 | 1.90 | 0.09 |
| (0.11) | (0.14) | (0.12) | | | (0.18) | | | | | | | |
| -0.08 | -0.23 | 1.09 | 0.03 | | | | | 5 | -785.23 | 1580.63 | 2.04 | 0.08 |
| (0.15) | (0.14) | (0.12) | (0.22) | | | | | | | | | |
| -0.06 | -0.23 | 1.09 | | -0.01 | | | | 5 | -785.24 | 1580.64 | 2.05 | 0.08 |
| (0.16) | (0.13) | (0.12) | | (0.22) | | | | | | | | |
| -0.12 | | 1.08 | 0.12 | | | | | 4 | -786.58 | 1581.27 | 2.68 | 0.06 |
| (0.15) | | (0.12) | (0.22) | | | | | | | | | |
| -0.05 | | 1.08 | | -0.04 | | | | 4 | -786.71 | 1581.53 | 2.94 | 0.05 |
| (0.16) | | (0.12) | | (0.22) | | | | | | | | |
| -0.10 | -0.12 | 1.09 | 0.04 | | | -0.15 | | 6 | -785.09 | 1582.41 | 3.82 | 0.03 |
| (0.15) | (0.24) | (0.12) | (0.22) | | | (0.29) | | | | | | |
| -0.08 | -0.22 | 1.08 | 0.03 | | -0.07 | | | 6 | -785.15 | 1582.54 | 3.95 | 0.03 |
| (0.15) | (0.14) | (0.12) | (0.22) | | (0.18) | | | | | | | |
| -0.06 | -0.22 | 1.08 | | -0.00 | -0.07 | | | 6 | -785.16 | 1582.56 | 3.97 | 0.03 |
| (0.16) | (0.14) | (0.12) | | (0.22) | (0.18) | | | | | | | |
| -0.06 | -0.19 | 1.09 | | -0.01 | | | -0.09 | 6 | -785.19 | 1582.60 | 4.01 | 0.03 |
| (0.16) | (0.19) | (0.12) | | (0.22) | | | (0.27) | | | | | |
| -0.07 | -0.23 | 1.09 | 0.03 | -0.01 | | | | 6 | -785.23 | 1582.69 | 4.10 | 0.03 |
| (0.19) | (0.14) | (0.12) | (0.22) | (0.22) | | | | | | | | |
| -0.10 | | 1.08 | 0.12 | -0.04 | | | | 5 | -786.57 | 1583.30 | 4.71 | 0.02 |
| (0.19) | | (0.12) | (0.22) | (0.22) | | | | | | | | |
| -0.10 | -0.12 | 1.08 | 0.04 | | -0.06 | -0.15 | | 7 | -785.02 | 1584.35 | 5.77 | 0.01 |
| (0.15) | (0.24) | (0.12) | (0.22) | | (0.18) | (0.30) | | | | | | |
| -0.09 | -0.12 | 1.09 | 0.04 | -0.01 | | -0.16 | | 7 | -785.09 | 1584.49 | 5.90 | 0.01 |
| (0.19) | (0.24) | (0.12) | (0.22) | (0.22) | | (0.30) | | | | | | |
| -0.06 | -0.17 | 1.09 | | -0.00 | -0.08 | | -0.10 | 7 | -785.09 | 1584.49 | 5.90 | 0.01 |
| (0.16) | (0.20) | (0.12) | | (0.22) | (0.18) | | (0.27) | | | | | |
| -0.08 | -0.22 | 1.08 | 0.03 | -0.01 | -0.07 | | | 7 | -785.15 | 1584.61 | 6.03 | 0.01 |
| (0.19) | (0.14) | (0.12) | (0.22) | (0.22) | (0.18) | | | | | | | |
| -0.07 | -0.18 | 1.09 | 0.03 | -0.01 | | | -0.09 | 7 | -785.18 | 1584.66 | 6.07 | 0.01 |
| (0.19) | (0.19) | (0.12) | (0.22) | (0.22) | | | (0.27) | | | | | |
| -0.09 | -0.12 | 1.08 | 0.04 | -0.01 | -0.06 | -0.15 | | 8 | -785.02 | 1586.44 | 7.85 | 0.00 |

| | | | | | | | | | | | | |
|---|---|---|---|---|---|---|---|---|---|---|---|---|
| (0.19) | (0.24) | (0.12) | (0.22) | (0.22) | (0.18) | (0.30) | | | | | | |
| -0.09 | -0.09 | 1.09 | 0.04 | -0.01 | | -0.15 | -0.08 | 8 | -785.05 | 1586.50 | 7.91 | 0.00 |
| (0.19) | (0.27) | (0.12) | (0.22) | (0.22) | | (0.30) | (0.27) | | | | | |
| -0.07 | -0.17 | 1.09 | 0.03 | -0.00 | -0.08 | | -0.10 | 8 | -785.08 | 1586.56 | 7.97 | 0.00 |
| (0.19) | (0.20) | (0.12) | (0.22) | (0.22) | (0.18) | | (0.27) | | | | | |
| **SHORT-TERM (ADULTS, N = 655 obs., n = 377 ind.)** | | | | | | | | | | | | |
| **I** | **FGM** | **Qcoh** | **Pop(CH)** | **Sex(M)** | **FGM:Qcoh** | **FGM:Pop(CH)** | **FGM:Sex(M)** | **df** | **Log-lik** | **AICc** | **Delta** | **Weight** |
| -0.01 | -0.32 | 0.26 | | | | | | 6 | -1268.53 | 2549.18 | 0.00 | 0.17 |
| (0.18) | (0.07) | (0.14) | | | | | | | | | | |
| -0.00 | -0.32 | 0.25 | | | 0.10 | | | 7 | -1267.99 | 2550.16 | 0.98 | 0.10 |
| (0.18) | (0.07) | (0.14) | | | (0.10) | | | | | | | |
| **-0.02** | **-0.32** | | | | | | | **5** | **-1270.38** | **2550.85** | **1.67** | **0.07** |
| **(0.18)** | **(0.07)** | | | | | | | | | | | |
| -0.07 | -0.31 | 0.26 | 0.14 | | | | | 7 | -1268.35 | 2550.87 | 1.69 | 0.07 |
| (0.21) | (0.07) | (0.14) | (0.23) | | | | | | | | | |
| -0.01 | -0.40 | 0.25 | | 0.01 | | | 0.20 | 8 | -1267.39 | 2551.01 | 1.83 | 0.07 |
| (0.20) | (0.09) | (0.14) | | (0.21) | | | (0.14) | | | | | |
| -0.01 | -0.32 | 0.26 | | 0.01 | | | | 7 | -1268.52 | 2551.22 | 2.04 | 0.06 |
| (0.20) | (0.07) | (0.14) | | (0.21) | | | | | | | | |
| -0.06 | -0.31 | 0.24 | 0.13 | | 0.10 | | | 8 | -1267.84 | 2551.91 | 2.73 | 0.04 |
| (0.21) | (0.07) | (0.14) | (0.23) | | (0.10) | | | | | | | |
| -0.00 | -0.39 | 0.24 | | -0.00 | 0.10 | | 0.20 | 9 | -1266.89 | 2552.05 | 2.87 | 0.04 |
| (0.20) | (0.09) | (0.14) | | (0.21) | (0.10) | | (0.14) | | | | | |
| -0.00 | -0.32 | 0.25 | | 0.00 | 0.10 | | | 8 | -1267.99 | 2552.21 | 3.03 | 0.04 |
| (0.20) | (0.07) | (0.14) | | (0.21) | (0.10) | | | | | | | |
| -0.10 | -0.23 | 0.26 | 0.15 | | | -0.12 | | 8 | -1268.00 | 2552.23 | 3.05 | 0.04 |
| (0.22) | (0.12) | (0.14) | (0.23) | | | (0.15) | | | | | | |
| -0.01 | -0.41 | | | -0.01 | | | 0.22 | 7 | -1269.07 | 2552.32 | 3.14 | 0.04 |
| (0.21) | (0.08) | | | (0.21) | | | (0.14) | | | | | |
| -0.09 | -0.31 | | 0.16 | | | | | 6 | -1270.13 | 2552.40 | 3.22 | 0.03 |
| (0.22) | (0.07) | | (0.23) | | | | | | | | | |
| -0.08 | -0.39 | 0.25 | 0.14 | 0.01 | | | 0.21 | 9 | -1267.20 | 2552.67 | 3.49 | 0.03 |
| (0.23) | (0.09) | (0.14) | (0.23) | (0.21) | | | (0.14) | | | | | |
| -0.01 | -0.32 | | | -0.01 | | | | 6 | -1270.38 | 2552.89 | 3.71 | 0.03 |
| (0.21) | (0.07) | | | (0.21) | | | | | | | | |
| -0.08 | -0.31 | 0.26 | 0.14 | 0.02 | | | | 8 | -1268.35 | 2552.91 | 3.73 | 0.03 |
| (0.23) | (0.07) | (0.14) | (0.23) | (0.21) | | | | | | | | |
| -0.09 | -0.24 | 0.25 | 0.14 | | 0.08 | -0.09 | | 9 | -1267.64 | 2553.56 | 4.38 | 0.02 |
| (0.21) | (0.12) | (0.14) | (0.23) | | (0.10) | (0.15) | | | | | | |
| -0.07 | -0.39 | 0.23 | 0.13 | 0.00 | 0.09 | | 0.20 | 10 | -1266.72 | 2553.78 | 4.60 | 0.02 |
| (0.23) | (0.09) | (0.14) | (0.23) | (0.21) | (0.10) | | (0.14) | | | | | |
| -0.12 | -0.23 | | 0.18 | | | -0.11 | | 7 | -1269.82 | 2553.82 | 4.64 | 0.02 |
| (0.22) | (0.12) | | (0.23) | | | (0.15) | | | | | | |
| -0.09 | -0.40 | | 0.17 | -0.01 | | | 0.22 | 8 | -1268.81 | 2553.84 | 4.66 | 0.02 |
| (0.24) | (0.09) | | (0.23) | (0.21) | | | (0.14) | | | | | |
| -0.07 | -0.31 | 0.25 | 0.13 | 0.00 | 0.10 | | | 9 | -1267.84 | 2553.96 | 4.78 | 0.02 |

| | | | | | | | | | | | | |
|---|---|---|---|---|---|---|---|---|---|---|---|---|
| (0.23) | (0.07) | (0.14) | (0.23) | (0.21) | (0.10) | | | | | | | |
| -0.11 | -0.23 | 0.26 | 0.15 | 0.02 | | -0.12 | | 9 | -1268.00 | 2554.28 | 5.10 | 0.01 |
| (0.23) | (0.12) | (0.14) | (0.23) | (0.21) | | (0.15) | | | | | | |
| -0.10 | -0.32 | 0.25 | 0.16 | 0.01 | | -0.09 | 0.19 | 10 | -1267.01 | 2554.37 | 5.19 | 0.01 |
| (0.24) | (0.14) | (0.14) | (0.23) | (0.21) | | (0.15) | (0.14) | | | | | |
| -0.09 | -0.31 | | 0.16 | -0.01 | | | | 7 | -1270.13 | 2554.44 | 5.26 | 0.01 |
| (0.24) | (0.07) | | (0.23) | (0.21) | | | | | | | | |
| -0.11 | -0.34 | | 0.18 | -0.01 | | -0.08 | 0.21 | 9 | -1268.66 | 2555.60 | 6.42 | 0.01 |
| (0.24) | (0.14) | | (0.23) | (0.21) | | (0.15) | (0.14) | | | | | |
| -0.09 | -0.24 | 0.25 | 0.14 | 0.01 | 0.08 | -0.09 | | 10 | -1267.64 | 2555.62 | 6.44 | 0.01 |
| (0.23) | (0.12) | (0.14) | (0.23) | (0.21) | (0.10) | (0.15) | | | | | | |
| -0.08 | -0.34 | 0.24 | 0.14 | 0.00 | 0.09 | -0.06 | 0.19 | 11 | -1266.63 | 2555.68 | 6.50 | 0.01 |
| (0.24) | (0.14) | (0.14) | (0.23) | (0.21) | (0.10) | (0.15) | (0.14) | | | | | |
| -0.12 | -0.23 | | 0.18 | -0.01 | | -0.11 | | 8 | -1269.82 | 2555.87 | 6.69 | 0.01 |
| (0.24) | (0.12) | | (0.23) | (0.21) | | (0.15) | | | | | | |

Table S3. Linear mixed effects model (LMM) selection table for the medium-term relationship between the change of body mass between two consecutive years for early-growing (*i.e.* between the first and second years of life), late-growing (*i.e.* between 2 and 4 years old) and prime age (*i.e.* between 4 and 10 years old) individuals, and faecal glucocorticoid metabolites ('FGM') measured age *t* ('FGM*t*') or FGMs considered as the mean FGM value of the FGMs measured at age *t* and age *t+1* ('mean FGM'). We accounted for cohort quality ('Qcoh'), population (reference level is Chizé: CH, and is compared to Trois-Fontaines: TF), and sex (reference level is male: M, and is compared to females: F) and the initial body mass (*i.e.* measured year *t*, 'Mass'). Values give the parameter coefficient and values between brackets in grey rows are the standard-errors. 'I' is the Intercept, 'df' is the number of parameters, 'Log-lik' is the log-likelihood, 'delta' is the difference of AICc between the candidate model and the model having the lowest AICc, and 'weight' the AIC weight of each model. Retained model is in bold. Only models up to delta AICc = 7 are displayed.

| **MEDIUM-TERM (EARLY GROWTH, FGMt, N = 99)** | | | | | | | | | | | | | | |
|---|---|---|---|---|---|---|---|---|---|---|---|---|---|---|
| **I** | **FGM** | **Qcoh** | **Pop(CH)** | **Sex(M)** | **Mass** | **FGM:Qcoh** | **FGM:Pop(CH)** | **FGM:Sex(M)** | **FGM:Mass** | **df** | **Log-lik** | **AICc** | **Delta** | **Weight** |
| 0.08 | | | -1.03 | | -0.19 | | | | | 5 | -187.73 | 386.10 | 0.00 | 0.11 |
| (0.31) | | | (0.42) | | (0.08) | | | | | | | | | |
| 0.14 | -0.24 | | -1.16 | | -0.20 | | | | | 6 | -187.09 | 387.09 | 0.99 | 0.07 |
| (0.30) | (0.21) | | (0.43) | | (0.08) | | | | | | | | | |
| 0.41 | | -0.30 | -1.75 | | -0.16 | | | | | 6 | -187.22 | 387.35 | 1.24 | 0.06 |
| (0.45) | | (0.29) | (0.83) | | (0.09) | | | | | | | | | |
| 0.05 | | | -1.03 | 0.07 | -0.19 | | | | | 6 | -187.70 | 388.32 | 2.22 | 0.04 |
| (0.34) | | | (0.42) | (0.32) | (0.08) | | | | | | | | | |
| 0.48 | | -0.50 | -1.90 | | | | | | | 5 | -188.87 | 388.39 | 2.29 | 0.04 |
| (0.46) | | (0.28) | (0.86) | | | | | | | | | | | |
| 0.20 | -0.50 | | -1.20 | | -0.21 | | 0.41 | | | 7 | -186.64 | 388.52 | 2.41 | 0.03 |
| (0.30) | (0.34) | | (0.43) | | (0.08) | | (0.43) | | | | | | | |
| 0.42 | -0.22 | -0.26 | -1.78 | | -0.17 | | | | | 7 | -186.69 | 388.61 | 2.50 | 0.03 |
| (0.43) | (0.21) | (0.29) | (0.81) | | (0.09) | | | | | | | | | |
| 0.55 | -0.41 | -0.33 | -1.98 | | -0.15 | -0.38 | | | 0.25 | 9 | -184.40 | 388.82 | 2.72 | 0.03 |
| (0.44) | (0.22) | (0.29) | (0.82) | | (0.09) | (0.18) | | | (0.14) | | | | | |
| **-0.35** | | | | | | | | | | **3** | **-191.34** | **388.94** | **2.83** | **0.03** |
| **(0.29)** | | | | | | | | | | | | | | |

| | | | | | | | | | | | | | | |
|---|---|---|---|---|---|---|---|---|---|---|---|---|---|---|
| 0.12 | -0.24 | | -1.16 | | -0.20 | | | | 0.06 | 7 | -186.95 | 389.13 | 3.02 | 0.02 |
| (0.31) | (0.21) | | (0.43) | | (0.08) | | | | (0.11) | | | | | |
| -0.33 | | | | | -0.10 | | | | | 4 | -190.37 | 389.17 | 3.06 | 0.02 |
| (0.29) | | | | | (0.07) | | | | | | | | | |
| -0.14 | | | -0.54 | | | | | | | 4 | -190.42 | 389.26 | 3.16 | 0.02 |
| (0.31) | | | (0.39) | | | | | | | | | | | |
| -0.36 | | 0.23 | | | -0.17 | | | | | 5 | -189.32 | 389.28 | 3.18 | 0.02 |
| (0.28) | | (0.15) | | | (0.09) | | | | | | | | | |
| 0.09 | -0.24 | | -1.18 | 0.10 | -0.21 | | | | | 7 | -187.03 | 389.30 | 3.19 | 0.02 |
| (0.34) | (0.21) | | (0.43) | (0.32) | (0.08) | | | | | | | | | |
| 0.20 | -0.72 | | -1.22 | | -0.20 | | 0.74 | | 0.16 | 8 | -185.90 | 389.40 | 3.30 | 0.02 |
| (0.31) | (0.39) | | (0.43) | | (0.08) | | (0.51) | | (0.13) | | | | | |
| 0.47 | -0.30 | -0.25 | -1.79 | | -0.17 | -0.17 | | | | 8 | -185.99 | 389.58 | 3.47 | 0.02 |
| (0.42) | (0.21) | (0.27) | (0.79) | | (0.09) | (0.14) | | | | | | | | |
| 0.39 | | -0.29 | -1.74 | 0.03 | -0.16 | | | | | 7 | -187.21 | 389.66 | 3.55 | 0.02 |
| (0.49) | | (0.29) | (0.84) | (0.32) | (0.09) | | | | | | | | | |
| 0.46 | -0.48 | -0.24 | -1.79 | | -0.18 | | 0.41 | | | 8 | -186.26 | 390.12 | 4.01 | 0.01 |
| (0.42) | (0.34) | (0.28) | (0.80) | | (0.09) | | (0.43) | | | | | | | |
| 0.48 | -0.15 | -0.48 | -1.92 | | | | | | | 6 | -188.62 | 390.15 | 4.04 | 0.01 |
| (0.45) | (0.21) | (0.28) | (0.84) | | | | | | | | | | | |
| 0.10 | -0.47 | | -1.22 | 0.10 | -0.20 | | | 0.50 | | 8 | -186.28 | 390.16 | 4.05 | 0.01 |
| (0.33) | (0.27) | | (0.43) | (0.31) | (0.08) | | | (0.41) | | | | | | |
| 0.42 | -0.22 | -0.28 | -1.83 | | -0.17 | | | | 0.07 | 8 | -186.50 | 390.60 | 4.50 | 0.01 |
| (0.44) | (0.21) | (0.29) | (0.83) | | (0.09) | | | | (0.11) | | | | | |
| 0.49 | | -0.50 | -1.91 | -0.02 | | | | | | 6 | -188.87 | 390.66 | 4.55 | 0.01 |
| (0.50) | | (0.28) | (0.86) | (0.32) | | | | | | | | | | |
| -0.36 | -0.19 | 0.27 | | | -0.19 | | | | | 6 | -188.91 | 390.73 | 4.63 | 0.01 |
| (0.27) | (0.21) | (0.16) | | | (0.09) | | | | | | | | | |
| -0.11 | -0.17 | | -0.62 | | | | | | | 5 | -190.09 | 390.82 | 4.71 | 0.01 |
| (0.31) | (0.21) | | (0.39) | | | | | | | | | | | |
| 0.16 | -0.50 | | -1.21 | 0.07 | -0.21 | | 0.40 | | | 8 | -186.62 | 390.84 | 4.73 | 0.01 |
| (0.34) | (0.34) | | (0.43) | (0.32) | (0.08) | | (0.44) | | | | | | | |
| 0.51 | -0.71 | -0.28 | -1.91 | | -0.17 | | 0.75 | | 0.17 | 9 | -185.42 | 390.86 | 4.75 | 0.01 |
| (0.44) | (0.39) | (0.29) | (0.82) | | (0.09) | | (0.51) | | (0.13) | | | | | |
| -0.35 | -0.10 | | | | | | | | | 4 | -191.23 | 390.89 | 4.79 | 0.01 |
| (0.29) | (0.21) | | | | | | | | | | | | | |
| -0.36 | | 0.06 | | | | | | | | 4 | -191.25 | 390.92 | 4.81 | 0.01 |
| (0.29) | | (0.13) | | | | | | | | | | | | |
| 0.37 | -0.22 | -0.25 | -1.77 | 0.07 | -0.18 | | | | | 8 | -186.67 | 390.93 | 4.83 | 0.01 |
| (0.47) | (0.21) | (0.29) | (0.82) | (0.32) | (0.09) | | | | | | | | | |
| -0.36 | | | | 0.03 | | | | | | 4 | -191.34 | 391.10 | 5.00 | 0.01 |
| (0.33) | | | | (0.32) | | | | | | | | | | |
| 0.53 | -0.24 | -0.47 | -1.92 | | | -0.17 | | | | 7 | -187.94 | 391.12 | 5.02 | 0.01 |
| (0.44) | (0.22) | (0.27) | (0.82) | | | (0.14) | | | | | | | | |
| -0.33 | -0.10 | | | | -0.10 | | | | | 5 | -190.26 | 391.17 | 5.07 | 0.01 |
| (0.29) | (0.21) | | | | (0.07) | | | | | | | | | |

| | | | | | | | | | | | | | | |
|---|---|---|---|---|---|---|---|---|---|---|---|---|---|---|
| 0.50 | -0.41 | -0.32 | -1.96 | 0.08 | -0.16 | -0.38 | | | 0.26 | 10 | -184.37 | 391.24 | 5.13 | 0.01 |
| (0.48) | (0.22) | (0.29) | (0.82) | (0.31) | (0.09) | (0.18) | | | (0.14) | | | | | |
| 0.55 | -0.36 | -0.33 | -1.98 | | -0.15 | -0.40 | -0.09 | | 0.25 | 10 | -184.39 | 391.29 | 5.18 | 0.01 |
| (0.44) | (0.45) | (0.29) | (0.82) | | (0.09) | (0.28) | (0.77) | | (0.14) | | | | | |
| 0.05 | -0.25 | | -1.17 | 0.13 | -0.20 | | | | 0.07 | 8 | -186.86 | 391.32 | 5.22 | 0.01 |
| (0.34) | (0.21) | | (0.43) | (0.32) | (0.08) | | | | (0.11) | | | | | |
| -0.35 | | | | 0.04 | -0.10 | | | | | 5 | -190.36 | 391.37 | 5.27 | 0.01 |
| (0.34) | | | | (0.32) | (0.07) | | | | | | | | | |
| -0.15 | | | -0.54 | 0.04 | | | | | | 5 | -190.41 | 391.47 | 5.37 | 0.01 |
| (0.35) | | | (0.39) | (0.32) | | | | | | | | | | |
| -0.40 | | 0.23 | | 0.08 | -0.18 | | | | | 6 | -189.29 | 391.49 | 5.38 | 0.01 |
| (0.32) | | (0.15) | | (0.32) | (0.09) | | | | | | | | | |
| 0.14 | -0.73 | | -1.23 | 0.12 | -0.21 | | 0.74 | | 0.16 | 9 | -185.83 | 391.69 | 5.59 | 0.01 |
| (0.34) | (0.39) | | (0.43) | (0.32) | (0.08) | | (0.51) | | (0.13) | | | | | |
| 0.17 | -0.71 | | -1.24 | 0.07 | -0.20 | | 0.38 | 0.49 | | 9 | -185.89 | 391.81 | 5.71 | 0.01 |
| (0.33) | (0.38) | | (0.43) | (0.32) | (0.08) | | (0.43) | (0.41) | | | | | | |
| -0.32 | -0.37 | 0.26 | | | -0.17 | -0.34 | | | 0.22 | 8 | -187.14 | 391.89 | 5.78 | 0.01 |
| (0.28) | (0.23) | (0.16) | | | (0.09) | (0.19) | | | (0.14) | | | | | |
| -0.31 | -0.26 | 0.26 | | | -0.19 | -0.16 | | | | 7 | -188.33 | 391.89 | 5.78 | 0.01 |
| (0.27) | (0.22) | (0.16) | | | (0.09) | (0.15) | | | | | | | | |
| 0.52 | -0.37 | -0.47 | -1.92 | | | | 0.34 | | | 7 | -188.33 | 391.89 | 5.78 | 0.01 |
| (0.44) | (0.34) | (0.27) | (0.83) | | | | (0.44) | | | | | | | |
| 0.06 | -0.50 | | -1.22 | 0.14 | -0.19 | | | 0.56 | 0.09 | 9 | -185.94 | 391.91 | 5.81 | 0.01 |
| (0.34) | (0.28) | | (0.43) | (0.32) | (0.08) | | | (0.41) | (0.11) | | | | | |
| 0.37 | -0.45 | -0.23 | -1.77 | 0.06 | -0.17 | | | 0.49 | | 9 | -185.95 | 391.92 | 5.81 | 0.01 |
| (0.46) | (0.27) | (0.28) | (0.80) | (0.32) | (0.09) | | | (0.41) | | | | | | |
| 0.15 | -1.03 | | -1.29 | 0.12 | -0.19 | | 0.79 | 0.60 | 0.20 | 10 | -184.74 | 391.98 | 5.88 | 0.01 |
| (0.34) | (0.44) | | (0.43) | (0.31) | (0.08) | | (0.51) | (0.41) | (0.13) | | | | | |
| 0.47 | -0.26 | -0.25 | -1.79 | | -0.17 | -0.20 | -0.09 | | | 9 | -185.98 | 391.99 | 5.89 | 0.01 |
| (0.42) | (0.45) | (0.28) | (0.79) | | (0.09) | (0.26) | (0.79) | | | | | | | |
| 0.46 | -0.30 | -0.25 | -1.79 | 0.02 | -0.18 | -0.17 | | | | 9 | -185.99 | 392.00 | 5.89 | 0.01 |
| (0.46) | (0.21) | (0.28) | (0.79) | (0.32) | (0.09) | (0.14) | | | | | | | | |
| 0.48 | -0.65 | -0.30 | -1.97 | 0.09 | -0.15 | -0.37 | | 0.53 | 0.28 | 11 | -183.52 | 392.07 | 5.96 | 0.01 |
| (0.47) | (0.29) | (0.28) | (0.81) | (0.31) | (0.09) | (0.18) | | (0.40) | (0.14) | | | | | |
| 0.48 | -0.15 | -0.48 | -1.92 | 0.00 | | | | | | 7 | -188.62 | 392.46 | 6.36 | 0.00 |
| (0.49) | (0.21) | (0.28) | (0.85) | (0.32) | | | | | | | | | | |
| 0.44 | -0.48 | -0.24 | -1.78 | 0.03 | -0.18 | | 0.40 | | | 9 | -186.25 | 392.53 | 6.43 | 0.00 |
| (0.47) | (0.34) | (0.28) | (0.80) | (0.32) | (0.09) | | (0.43) | | | | | | | |
| -0.06 | -0.38 | | -0.63 | | | | 0.33 | | | 6 | -189.82 | 392.55 | 6.44 | 0.00 |
| (0.31) | (0.35) | | (0.39) | | | | (0.45) | | | | | | | |
| -0.36 | -0.13 | 0.08 | | | | | | | | 5 | -191.07 | 392.79 | 6.68 | 0.00 |
| (0.28) | (0.21) | (0.13) | | | | | | | | | | | | |
| -0.38 | -0.20 | 0.26 | | | -0.18 | | | | 0.06 | 7 | -188.78 | 392.80 | 6.69 | 0.00 |
| (0.28) | (0.21) | (0.16) | | | (0.09) | | | | (0.11) | | | | | |
| -0.42 | -0.20 | 0.27 | | 0.11 | -0.19 | | | | | 7 | -188.85 | 392.92 | 6.82 | 0.00 |
| (0.32) | (0.21) | (0.16) | | (0.32) | (0.09) | | | | | | | | | |

| | | | | | | | | | | | | | | |
|---|---|---|---|---|---|---|---|---|---|---|---|---|---|---|
| 0.36 | -0.23 | -0.27 | -1.81 | 0.10 | -0.17 | | | | 0.07 | 9 | -186.45 | 392.93 | 6.83 | 0.00 |
| (0.48) | (0.21) | (0.29) | (0.83) | (0.32) | (0.09) | | | | (0.11) | | | | | |
| -0.14 | -0.18 | | -0.62 | 0.06 | | | | | | 6 | -190.07 | 393.05 | 6.95 | 0.00 |
| (0.35) | (0.21) | | (0.39) | (0.32) | | | | | | | | | | |
| -0.37 | -0.10 | | | 0.04 | | | | | | 5 | -191.23 | 393.10 | 6.99 | 0.00 |
| (0.33) | (0.21) | | | (0.32) | | | | | | | | | | |
| **MEDIUM-TERM (EARLY GROWTH, mean FGM, N = 67)** | | | | | | | | | | | | | | |
| **I** | **FGM** | **Qcoh** | **Pop(CH)** | **Sex(M)** | **Mass** | **FGM:Qcoh** | **FGM:Pop(CH)** | **FGM:Sex(M)** | **FGM:Mass** | **df** | **Log-lik** | **AICc** | **Delta** | **Weight** |
| 0.32 | -0.85 | | -1.57 | | -0.27 | | | | | 6 | -127.12 | 267.65 | 0.00 | 0.10 |
| (0.34) | (0.34) | | (0.54) | | (0.10) | | | | | | | | | |
| 0.16 | -0.92 | | -1.75 | 0.46 | -0.28 | | | | | 7 | -126.48 | 268.87 | 1.22 | 0.06 |
| (0.37) | (0.34) | | (0.52) | (0.39) | (0.09) | | | | | | | | | |
| 0.75 | -0.98 | -0.37 | -2.43 | | -0.21 | -0.72 | | | 0.47 | 9 | -123.91 | 268.97 | 1.32 | 0.05 |
| (0.47) | (0.34) | (0.30) | (0.91) | | (0.10) | (0.31) | | | (0.20) | | | | | |
| 0.54 | -0.85 | -0.24 | -2.11 | | -0.23 | | | | | 7 | -126.77 | 269.45 | 1.80 | 0.04 |
| (0.43) | (0.34) | (0.28) | (0.85) | | (0.11) | | | | | | | | | |
| 0.40 | -1.07 | | -1.68 | | -0.29 | | 0.44 | | | 7 | -126.95 | 269.79 | 2.14 | 0.04 |
| (0.33) | (0.48) | | (0.53) | | (0.10) | | (0.70) | | | | | | | |
| 0.30 | -0.84 | | -1.53 | | -0.26 | | | | 0.07 | 7 | -127.01 | 269.92 | 2.27 | 0.03 |
| (0.35) | (0.34) | | (0.55) | | (0.10) | | | | (0.14) | | | | | |
| **-0.37** | | | | | | | | | | **3** | **-131.93** | **270.23** | **2.58** | **0.03** |
| **(0.38)** | | | | | | | | | | | | | | |
| 0.03 | | | -0.94 | | -0.21 | | | | | 5 | -129.73 | 270.45 | 2.80 | 0.03 |
| (0.39) | | | (0.55) | | (0.10) | | | | | | | | | |
| -0.37 | | | | | -0.12 | | | | | 4 | -130.93 | 270.51 | 2.87 | 0.02 |
| (0.37) | | | | | (0.09) | | | | | | | | | |
| -0.34 | -0.47 | | | | | | | | | 4 | -130.99 | 270.63 | 2.98 | 0.02 |
| (0.36) | (0.34) | | | | | | | | | | | | | |
| 0.54 | -1.03 | -0.33 | -2.43 | 0.40 | -0.22 | -0.72 | | | 0.48 | 10 | -123.35 | 270.63 | 2.98 | 0.02 |
| (0.49) | (0.33) | (0.29) | (0.88) | (0.38) | (0.10) | (0.31) | | | (0.20) | | | | | |
| -0.35 | -0.49 | | | | -0.13 | | | | | 5 | -129.89 | 270.77 | 3.12 | 0.02 |
| (0.36) | (0.33) | | | | (0.09) | | | | | | | | | |
| 0.57 | -0.90 | -0.21 | -2.09 | | -0.25 | -0.23 | | | | 8 | -126.23 | 270.95 | 3.30 | 0.02 |
| (0.41) | (0.34) | (0.27) | (0.82) | | (0.11) | (0.21) | | | | | | | | |
| 0.24 | -1.17 | | -1.88 | 0.48 | -0.30 | | 0.49 | | | 8 | -126.26 | 271.01 | 3.36 | 0.02 |
| (0.36) | (0.47) | | (0.51) | (0.39) | (0.10) | | (0.70) | | | | | | | |
| 0.34 | -0.90 | -0.18 | -2.10 | 0.40 | -0.24 | | | | | 8 | -126.28 | 271.05 | 3.40 | 0.02 |
| (0.47) | (0.34) | (0.27) | (0.82) | (0.40) | (0.11) | | | | | | | | | |
| 0.14 | -0.92 | | -1.72 | 0.48 | -0.27 | | | | 0.09 | 8 | -126.32 | 271.12 | 3.47 | 0.02 |
| (0.38) | (0.34) | | (0.54) | (0.40) | (0.09) | | | | (0.14) | | | | | |
| 0.53 | -0.67 | -0.52 | -2.10 | | | | | | | 6 | -128.89 | 271.19 | 3.54 | 0.02 |
| (0.49) | (0.35) | (0.28) | (0.94) | | | | | | | | | | | |
| 0.45 | -1.36 | | -1.73 | | -0.29 | | 1.00 | | 0.20 | 8 | -126.40 | 271.29 | 3.64 | 0.02 |
| (0.36) | (0.58) | | (0.56) | | (0.10) | | (0.90) | | (0.18) | | | | | |
| 0.63 | -0.62 | -0.35 | -2.23 | | -0.19 | -0.91 | -0.75 | | 0.47 | 10 | -123.75 | 271.42 | 3.77 | 0.02 |
| (0.51) | (0.69) | (0.31) | (0.98) | | (0.10) | (0.45) | (1.31) | | (0.20) | | | | | |

| | | | | | | | | | | | | | | |
|---|---|---|---|---|---|---|---|---|---|---|---|---|---|---|
| 0.16 | -0.94 | | -1.76 | 0.46 | -0.28 | | | 0.05 | | 8 | -126.48 | 271.44 | 3.80 | 0.02 |
| (0.37) | (0.44) | | (0.52) | (0.39) | (0.10) | | | (0.67) | | | | | | |
| -0.34 | -0.62 | 0.25 | | | -0.22 | | | | | 6 | -129.08 | 271.56 | 3.91 | 0.01 |
| (0.33) | (0.34) | (0.19) | | | (0.11) | | | | | | | | | |
| 0.57 | -0.83 | -0.29 | -2.20 | | -0.21 | | | | 0.10 | 8 | -126.54 | 271.57 | 3.92 | 0.01 |
| (0.47) | (0.34) | (0.30) | (0.92) | | (0.11) | | | | (0.14) | | | | | |
| 0.63 | -1.10 | -0.24 | -2.23 | | -0.25 | | 0.47 | | | 8 | -126.57 | 271.62 | 3.98 | 0.01 |
| (0.42) | (0.47) | (0.27) | (0.83) | | (0.11) | | (0.70) | | | | | | | |
| -0.11 | -0.59 | | -0.55 | | | | | | | 5 | -130.47 | 271.92 | 4.28 | 0.01 |
| (0.39) | (0.35) | | (0.49) | | | | | | | | | | | |
| 0.29 | -1.55 | | -2.00 | 0.56 | -0.31 | | 1.18 | | 0.23 | 9 | -125.49 | 272.14 | 4.49 | 0.01 |
| (0.38) | (0.57) | | (0.55) | (0.40) | (0.10) | | (0.90) | | (0.18) | | | | | |
| 0.33 | | -0.49 | -1.70 | | | | | | | 5 | -130.58 | 272.14 | 4.50 | 0.01 |
| (0.54) | | (0.32) | (1.01) | | | | | | | | | | | |
| 0.29 | | -0.27 | -1.59 | | -0.17 | | | | | 6 | -129.39 | 272.18 | 4.53 | 0.01 |
| (0.51) | | (0.33) | (0.97) | | (0.11) | | | | | | | | | |
| -0.25 | | | -0.28 | | | | | | | 4 | -131.77 | 272.19 | 4.55 | 0.01 |
| (0.41) | | | (0.49) | | | | | | | | | | | |
| -0.36 | | 0.15 | | | -0.18 | | | | | 5 | -130.63 | 272.24 | 4.59 | 0.01 |
| (0.36) | | (0.19) | | | (0.11) | | | | | | | | | |
| -0.47 | | | | 0.20 | | | | | | 4 | -131.81 | 272.26 | 4.61 | 0.01 |
| (0.43) | | | | (0.41) | | | | | | | | | | |
| -0.09 | | | -1.01 | 0.30 | -0.21 | | | | | 6 | -129.46 | 272.32 | 4.67 | 0.01 |
| (0.42) | | | (0.55) | (0.40) | (0.10) | | | | | | | | | |
| -0.27 | -0.75 | 0.24 | | | -0.21 | -0.64 | | | 0.39 | 8 | -126.92 | 272.33 | 4.68 | 0.01 |
| (0.35) | (0.34) | (0.19) | | | (0.11) | (0.31) | | | (0.20) | | | | | |
| -0.37 | | -0.04 | | | | | | | | 4 | -131.90 | 272.44 | 4.79 | 0.01 |
| (0.38) | | (0.16) | | | | | | | | | | | | |
| 0.83 | -1.48 | -0.37 | -2.62 | | -0.24 | | 1.23 | | 0.26 | 9 | -125.65 | 272.46 | 4.82 | 0.01 |
| (0.49) | (0.58) | (0.30) | (0.94) | | (0.11) | | (0.91) | | (0.18) | | | | | |
| -0.46 | -0.48 | | | 0.24 | | | | | | 5 | -130.82 | 272.62 | 4.97 | 0.01 |
| (0.41) | (0.34) | | | (0.41) | | | | | | | | | | |
| -0.47 | | | | 0.19 | -0.12 | | | | | 5 | -130.82 | 272.63 | 4.98 | 0.01 |
| (0.42) | | | | (0.41) | (0.09) | | | | | | | | | |
| 0.37 | -0.94 | -0.15 | -2.08 | 0.38 | -0.26 | -0.22 | | | | 9 | -125.80 | 272.75 | 5.10 | 0.01 |
| (0.46) | (0.34) | (0.26) | (0.79) | (0.40) | (0.11) | (0.21) | | | | | | | | |
| -0.36 | -0.49 | | | | -0.12 | | | | 0.09 | 6 | -129.71 | 272.82 | 5.17 | 0.01 |
| (0.38) | (0.33) | | | | (0.09) | | | | (0.14) | | | | | |
| -0.46 | -0.50 | | | 0.23 | -0.13 | | | | | 6 | -129.73 | 272.86 | 5.21 | 0.01 |
| (0.41) | (0.33) | | | (0.40) | (0.09) | | | | | | | | | |
| -0.34 | -0.47 | 0.01 | | | | | | | | 5 | -130.99 | 272.96 | 5.31 | 0.01 |
| (0.36) | (0.35) | (0.16) | | | | | | | | | | | | |
| -0.30 | -0.66 | 0.26 | | | -0.23 | -0.22 | | | | 7 | -128.58 | 273.05 | 5.40 | 0.01 |
| (0.32) | (0.34) | (0.19) | | | (0.11) | (0.22) | | | | | | | | |
| 0.55 | -0.71 | -0.50 | -2.07 | | | -0.17 | | | | 7 | -128.62 | 273.13 | 5.49 | 0.01 |
| (0.47) | (0.35) | (0.27) | (0.92) | | | (0.22) | | | | | | | | |

| | | | | | | | | | | | | | | |
|---|---|---|---|---|---|---|---|---|---|---|---|---|---|---|
| 0.36 | -0.89 | -0.24 | -2.20 | 0.42 | -0.23 | | | | 0.11 | 9 | -126.01 | 273.17 | 5.52 | 0.01 |
| (0.49) | (0.34) | (0.29) | (0.89) | (0.40) | (0.11) | | | | (0.14) | | | | | |
| 0.38 | -0.70 | -0.49 | -2.10 | 0.29 | | | | | | 7 | -128.64 | 273.18 | 5.53 | 0.01 |
| (0.52) | (0.35) | (0.28) | (0.92) | (0.40) | | | | | | | | | | |
| 0.42 | -1.17 | -0.17 | -2.23 | 0.42 | -0.27 | | 0.50 | | | 9 | -126.05 | 273.26 | 5.62 | 0.01 |
| (0.47) | (0.47) | (0.26) | (0.79) | (0.40) | (0.11) | | (0.70) | | | | | | | |
| 0.48 | -0.77 | -0.31 | -2.29 | 0.38 | -0.21 | -0.86 | -0.55 | | 0.48 | 11 | -123.27 | 273.34 | 5.69 | 0.01 |
| (0.52) | (0.69) | (0.30) | (0.95) | (0.38) | (0.10) | (0.45) | (1.31) | | (0.20) | | | | | |
| -0.51 | -0.65 | 0.29 | | 0.35 | -0.23 | | | | | 7 | -128.73 | 273.35 | 5.71 | 0.01 |
| (0.38) | (0.34) | (0.19) | | (0.40) | (0.11) | | | | | | | | | |
| 0.47 | -0.58 | -0.18 | -1.91 | | -0.23 | -0.40 | -0.67 | | | 9 | -126.12 | 273.39 | 5.75 | 0.01 |
| (0.46) | (0.71) | (0.28) | (0.89) | | (0.11) | (0.42) | (1.37) | | | | | | | |
| 0.57 | -0.92 | -0.34 | -2.45 | 0.39 | -0.22 | -0.74 | | -0.23 | 0.48 | 11 | -123.30 | 273.39 | 5.75 | 0.01 |
| (0.50) | (0.45) | (0.29) | (0.89) | (0.38) | (0.10) | (0.31) | | (0.68) | (0.20) | | | | | |
| 0.24 | -1.15 | | -1.88 | 0.47 | -0.30 | | 0.51 | -0.08 | | 9 | -126.26 | 273.67 | 6.02 | 0.01 |
| (0.37) | (0.50) | | (0.52) | (0.39) | (0.10) | | (0.74) | (0.71) | | | | | | |
| 0.55 | -0.71 | -0.52 | -2.12 | | | | 0.07 | | | 7 | -128.89 | 273.67 | 6.03 | 0.01 |
| (0.49) | (0.48) | (0.28) | (0.94) | | | | (0.71) | | | | | | | |
| -0.24 | -0.63 | | -0.64 | 0.34 | | | | | | 6 | -130.14 | 273.69 | 6.04 | 0.01 |
| (0.42) | (0.35) | | (0.49) | (0.41) | | | | | | | | | | |
| 0.34 | -0.89 | -0.18 | -2.11 | 0.40 | -0.24 | | | -0.03 | | 9 | -126.28 | 273.72 | 6.08 | 0.01 |
| (0.48) | (0.44) | (0.28) | (0.83) | (0.40) | (0.11) | | | (0.67) | | | | | | |
| 0.13 | -1.00 | | -1.74 | 0.49 | -0.27 | | | 0.18 | 0.10 | 9 | -126.29 | 273.74 | 6.09 | 0.00 |
| (0.37) | (0.47) | | (0.54) | (0.40) | (0.10) | | | (0.71) | (0.15) | | | | | |
| 0.61 | -1.61 | -0.31 | -2.66 | 0.48 | -0.26 | | 1.35 | | 0.29 | 10 | -124.94 | 273.81 | 6.16 | 0.00 |
| (0.50) | (0.58) | (0.28) | (0.90) | (0.39) | (0.11) | | (0.91) | | (0.19) | | | | | |
| -0.35 | -0.62 | 0.24 | | | -0.21 | | | | 0.07 | 7 | -128.96 | 273.82 | 6.17 | 0.00 |
| (0.35) | (0.34) | (0.19) | | | (0.11) | | | | (0.14) | | | | | |
| -0.35 | | | -0.33 | 0.24 | | | | | | 5 | -131.61 | 274.20 | 6.55 | 0.00 |
| (0.45) | | | (0.49) | (0.41) | | | | | | | | | | |
| 0.15 | | -0.24 | -1.58 | 0.26 | -0.18 | | | | | 7 | -129.19 | 274.27 | 6.62 | 0.00 |
| (0.55) | | (0.33) | (0.96) | (0.40) | (0.11) | | | | | | | | | |
| -0.49 | | 0.17 | | 0.25 | -0.18 | | | | | 6 | -130.44 | 274.28 | 6.63 | 0.00 |
| (0.41) | | (0.19) | | (0.41) | (0.11) | | | | | | | | | |
| -0.44 | -0.78 | 0.27 | | 0.34 | -0.21 | -0.64 | | | 0.40 | 9 | -126.56 | 274.28 | 6.64 | 0.00 |
| (0.40) | (0.34) | (0.19) | | (0.39) | (0.11) | (0.31) | | | (0.20) | | | | | |
| -0.12 | -0.52 | | -0.54 | | | | -0.14 | | | 6 | -130.45 | 274.31 | 6.66 | 0.00 |
| (0.39) | (0.49) | | (0.49) | | | | (0.72) | | | | | | | |
| 0.23 | | -0.48 | -1.69 | 0.20 | | | | | | 6 | -130.46 | 274.31 | 6.67 | 0.00 |
| (0.57) | | (0.31) | (1.01) | (0.41) | | | | | | | | | | |
| -0.46 | | -0.03 | | 0.19 | | | | | | 5 | -131.79 | 274.57 | 6.92 | 0.00 |
| (0.43) | | (0.16) | | (0.41) | | | | | | | | | | |
| **MEDIUM-TERM (LATE GROWTH, FGMt, N = 85)** | | | | | | | | | | | | | | |
| **I** | **FGM** | **Qcoh** | **Pop(CH)** | **Sex(M)** | **Mass** | **FGM:Qcoh** | **FGM:Pop(CH)** | **FGM:Sex(M)** | **FGM:Mass** | **df** | **Log-lik** | **AICc** | **Delta** | **Weight** |
| -0.54 | 0.35 | | | | -0.15 | | | | | 5 | -133.70 | 278.15 | 0.00 | 0.16 |
| (0.23) | (0.18) | | | | (0.05) | | | | | | | | | |

| | | | | | | | | | | | | | | |
|---|---|---|---|---|---|---|---|---|---|---|---|---|---|---|
| -0.52 | 0.26 | | | | -0.15 | | | | -0.08 | 6 | -133.10 | 279.28 | 1.12 | 0.09 |
| (0.23) | (0.20) | | | | (0.05) | | | | (0.08) | | | | | |
| **-0.55** | | | | | **-0.15** | | | | | **4** | **-135.49** | **279.47** | **1.32** | **0.08** |
| **(0.24)** | | | | | **(0.05)** | | | | | | | | | |
| -0.52 | 0.34 | | | -0.04 | -0.15 | | | | | 6 | -133.68 | 280.45 | 2.29 | 0.05 |
| (0.26) | (0.18) | | | (0.26) | (0.05) | | | | | | | | | |
| -0.56 | 0.36 | | 0.04 | | -0.15 | | | | | 6 | -133.69 | 280.46 | 2.30 | 0.05 |
| (0.29) | (0.22) | | (0.36) | | (0.06) | | | | | | | | | |
| -0.54 | 0.34 | 0.01 | | | -0.15 | | | | | 6 | -133.69 | 280.46 | 2.31 | 0.05 |
| (0.23) | (0.20) | (0.10) | | | (0.05) | | | | | | | | | |
| -0.41 | | | -0.30 | | -0.17 | | | | | 5 | -135.03 | 280.83 | 2.67 | 0.04 |
| (0.27) | | | (0.31) | | (0.06) | | | | | | | | | |
| -0.54 | | 0.09 | | | -0.17 | | | | | 5 | -135.04 | 280.84 | 2.68 | 0.04 |
| (0.23) | | (0.09) | | | (0.05) | | | | | | | | | |
| -0.50 | | | | -0.13 | -0.14 | | | | | 5 | -135.36 | 281.49 | 3.33 | 0.03 |
| (0.26) | | | | (0.26) | (0.05) | | | | | | | | | |
| -0.49 | 0.25 | | | -0.07 | -0.14 | | | | -0.09 | 7 | -133.07 | 281.59 | 3.44 | 0.03 |
| (0.25) | (0.20) | | | (0.26) | (0.05) | | | | (0.08) | | | | | |
| -0.55 | 0.28 | | 0.07 | | -0.14 | | | | -0.09 | 7 | -133.08 | 281.61 | 3.46 | 0.03 |
| (0.28) | (0.23) | | (0.36) | | (0.06) | | | | (0.08) | | | | | |
| -0.52 | 0.25 | 0.01 | | | -0.15 | | | | -0.08 | 7 | -133.09 | 281.64 | 3.49 | 0.03 |
| (0.23) | (0.22) | (0.10) | | | (0.05) | | | | (0.08) | | | | | |
| -0.62 | 0.52 | | 0.10 | | -0.14 | | -0.23 | | | 7 | -133.56 | 282.58 | 4.43 | 0.02 |
| (0.31) | (0.38) | | (0.37) | | (0.06) | | (0.46) | | | | | | | |
| -0.67 | 0.37 | 0.08 | 0.29 | | -0.14 | | | | | 7 | -133.62 | 282.69 | 4.53 | 0.02 |
| (0.39) | (0.22) | (0.20) | (0.69) | | (0.06) | | | | | | | | | |
| -0.54 | 0.36 | | 0.06 | -0.05 | -0.14 | | | | | 7 | -133.67 | 282.80 | 4.65 | 0.02 |
| (0.30) | (0.22) | | (0.36) | (0.27) | (0.06) | | | | | | | | | |
| -0.52 | 0.34 | 0.01 | | -0.04 | -0.15 | | | | | 7 | -133.68 | 282.82 | 4.67 | 0.02 |
| (0.26) | (0.21) | (0.11) | | (0.27) | (0.06) | | | | | | | | | |
| -0.52 | 0.34 | | | -0.04 | -0.15 | | | 0.00 | | 7 | -133.68 | 282.82 | 4.67 | 0.02 |
| (0.26) | (0.23) | | | (0.27) | (0.05) | | | (0.37) | | | | | | |
| -0.53 | 0.33 | 0.01 | | | -0.15 | -0.02 | | | | 7 | -133.69 | 282.83 | 4.67 | 0.02 |
| (0.25) | (0.22) | (0.11) | | | (0.05) | (0.14) | | | | | | | | |
| -0.39 | | | -0.28 | -0.08 | -0.17 | | | | | 6 | -135.00 | 283.07 | 4.92 | 0.01 |
| (0.28) | | | (0.31) | (0.27) | (0.06) | | | | | | | | | |
| -0.52 | | 0.08 | | -0.07 | -0.17 | | | | | 6 | -135.01 | 283.10 | 4.94 | 0.01 |
| (0.26) | | (0.10) | | (0.27) | (0.06) | | | | | | | | | |
| -0.47 | | 0.05 | -0.16 | | -0.17 | | | | | 6 | -135.01 | 283.10 | 4.95 | 0.01 |
| (0.37) | | (0.20) | (0.66) | | (0.06) | | | | | | | | | |
| -0.65 | 0.52 | | 0.16 | | -0.12 | | -0.35 | | -0.10 | 8 | -132.79 | 283.48 | 5.32 | 0.01 |
| (0.31) | (0.38) | | (0.37) | | (0.06) | | (0.46) | | (0.08) | | | | | |
| -0.72 | 0.29 | 0.12 | 0.45 | | -0.13 | | | | -0.09 | 8 | -132.91 | 283.71 | 5.56 | 0.01 |
| (0.39) | (0.22) | (0.20) | (0.70) | | (0.06) | | | | (0.08) | | | | | |
| -0.53 | 0.28 | | 0.09 | -0.08 | -0.13 | | | | -0.09 | 8 | -133.04 | 283.97 | 5.82 | 0.01 |
| (0.29) | (0.23) | | (0.36) | (0.27) | (0.06) | | | | (0.08) | | | | | |

| | | | | | | | | | | | | | | |
|---|---|---|---|---|---|---|---|---|---|---|---|---|---|---|
| -0.49 | 0.23 | | | -0.06 | -0.14 | | | 0.06 | -0.09 | 8 | -133.06 | 284.01 | 5.85 | 0.01 |
| (0.25) | (0.25) | | | (0.27) | (0.05) | | | (0.37) | (0.08) | | | | | |
| -0.49 | 0.25 | 0.01 | | -0.06 | -0.14 | | | | -0.09 | 8 | -133.07 | 284.03 | 5.87 | 0.01 |
| (0.25) | (0.22) | (0.11) | | (0.27) | (0.06) | | | | (0.08) | | | | | |
| -0.51 | 0.25 | 0.01 | | | -0.15 | -0.01 | | | -0.08 | 8 | -133.09 | 284.08 | 5.92 | 0.01 |
| (0.24) | (0.23) | (0.10) | | | (0.05) | (0.14) | | | (0.08) | | | | | |
| -0.80 | 0.52 | | 0.56 | | | | | | | 5 | -136.84 | 284.44 | 6.28 | 0.01 |
| (0.30) | (0.21) | | (0.31) | | | | | | | | | | | |
| -0.74 | 0.53 | 0.08 | 0.35 | | -0.13 | | -0.23 | | | 8 | -133.49 | 284.87 | 6.71 | 0.01 |
| (0.41) | (0.38) | (0.20) | (0.69) | | (0.06) | | (0.46) | | | | | | | |
| -0.61 | 0.51 | | 0.10 | -0.04 | -0.13 | | -0.23 | | | 8 | -133.55 | 285.00 | 6.84 | 0.01 |
| (0.33) | (0.38) | | (0.37) | (0.27) | (0.06) | | (0.46) | | | | | | | |
| -0.66 | 0.37 | 0.08 | 0.29 | -0.03 | -0.14 | | | | | 8 | -133.61 | 285.11 | 6.96 | 0.00 |
| (0.41) | (0.22) | (0.20) | (0.69) | (0.27) | (0.06) | | | | | | | | | |
| -0.66 | 0.36 | 0.08 | 0.28 | | -0.14 | -0.01 | | | | 8 | -133.61 | 285.12 | 6.97 | 0.00 |
| (0.41) | (0.24) | (0.20) | (0.70) | | (0.06) | (0.15) | | | | | | | | |
| **MEDIUM-TERM (LATE GROWTH, mean FGM, N = 63)** | | | | | | | | | | | | | | |
| **I** | **FGM** | **Qcoh** | **Pop(CH)** | **Sex(M)** | **Mass** | **FGM:Qcoh** | **FGM:Pop(CH)** | **FGM:Sex(M)** | **FGM:Mass** | **df** | **Log-lik** | **AICc** | **Delta** | **Weight** |
| **-0.47** | | | | | **-0.17** | | | | | **4** | **-97.55** | **203.78** | **0.00** | **0.16** |
| **(0.24)** | | | | | **(0.05)** | | | | | | | | | |
| -0.39 | -0.10 | | | | -0.12 | | | | -0.27 | 6 | -95.24 | 203.98 | 0.20 | 0.14 |
| (0.23) | (0.27) | | | | (0.06) | | | | (0.13) | | | | | |
| -0.46 | 0.16 | | | | -0.17 | | | | | 5 | -97.33 | 205.71 | 1.93 | 0.06 |
| (0.24) | (0.25) | | | | (0.05) | | | | | | | | | |
| -0.56 | 0.04 | | 0.35 | | -0.10 | | | | -0.27 | 7 | -94.94 | 205.92 | 2.14 | 0.05 |
| (0.32) | (0.32) | | (0.42) | | (0.06) | | | | (0.13) | | | | | |
| -0.43 | | | | -0.09 | -0.16 | | | | | 5 | -97.50 | 206.06 | 2.28 | 0.05 |
| (0.27) | | | | (0.31) | (0.05) | | | | | | | | | |
| -0.47 | | 0.03 | | | -0.17 | | | | | 5 | -97.51 | 206.08 | 2.30 | 0.05 |
| (0.24) | | (0.11) | | | (0.06) | | | | | | | | | |
| -0.47 | | | 0.01 | | -0.17 | | | | | 5 | -97.54 | 206.14 | 2.36 | 0.05 |
| (0.29) | | | (0.34) | | (0.06) | | | | | | | | | |
| -0.39 | -0.02 | -0.07 | | | -0.11 | | | | -0.27 | 7 | -95.13 | 206.30 | 2.52 | 0.04 |
| (0.24) | (0.32) | (0.14) | | | (0.06) | | | | (0.13) | | | | | |
| -0.36 | -0.12 | | | -0.09 | -0.12 | | | | -0.27 | 7 | -95.20 | 206.43 | 2.65 | 0.04 |
| (0.26) | (0.28) | | | (0.31) | (0.06) | | | | (0.13) | | | | | |
| -0.60 | 0.28 | | 0.27 | | -0.15 | | | | | 6 | -97.15 | 207.81 | 4.03 | 0.02 |
| (0.32) | (0.31) | | (0.44) | | (0.06) | | | | | | | | | |
| -0.88 | | 0.28 | 0.85 | | -0.16 | | | | | 6 | -97.19 | 207.88 | 4.10 | 0.02 |
| (0.52) | | (0.31) | (0.95) | | (0.06) | | | | | | | | | |
| -0.46 | 0.21 | -0.03 | | | -0.17 | | | | | 6 | -97.31 | 208.11 | 4.33 | 0.02 |
| (0.24) | (0.32) | (0.14) | | | (0.06) | | | | | | | | | |
| -0.45 | 0.15 | | | -0.05 | -0.17 | | | | | 6 | -97.32 | 208.14 | 4.36 | 0.02 |
| (0.26) | (0.25) | | | (0.32) | (0.05) | | | | | | | | | |
| -0.79 | 0.03 | 0.16 | 0.81 | | -0.10 | | | | -0.26 | 8 | -94.81 | 208.29 | 4.51 | 0.02 |
| (0.51) | (0.32) | (0.31) | (0.93) | | (0.06) | | | | (0.13) | | | | | |

| | | | | | | | | | | | | | | |
|---|---|---|---|---|---|---|---|---|---|---|---|---|---|---|
| -0.53 | 0.03 | | 0.37 | -0.12 | -0.09 | | | | -0.28 | 8 | -94.87 | 208.40 | 4.62 | 0.02 |
| (0.33) | (0.32) | | (0.43) | (0.31) | (0.07) | | | | (0.13) | | | | | |
| -0.30 | -0.15 | -0.03 | | | -0.12 | -0.15 | | | -0.26 | 8 | -94.90 | 208.46 | 4.68 | 0.02 |
| (0.27) | (0.38) | (0.15) | | | (0.07) | (0.22) | | | (0.13) | | | | | |
| -0.44 | | 0.02 | | -0.08 | -0.17 | | | | | 6 | -97.48 | 208.47 | 4.69 | 0.02 |
| (0.27) | | (0.12) | | (0.32) | (0.06) | | | | | | | | | |
| -0.38 | -0.00 | | | -0.14 | -0.11 | | | -0.39 | -0.28 | 8 | -94.91 | 208.48 | 4.70 | 0.02 |
| (0.26) | (0.32) | | | (0.31) | (0.06) | | | (0.50) | (0.13) | | | | | |
| -0.45 | | | 0.04 | -0.10 | -0.16 | | | | | 6 | -97.50 | 208.49 | 4.71 | 0.01 |
| (0.30) | | | (0.35) | (0.32) | (0.06) | | | | | | | | | |
| -0.63 | 0.20 | | 0.41 | | -0.09 | | -0.19 | | -0.28 | 8 | -94.92 | 208.50 | 4.72 | 0.01 |
| (0.43) | (0.77) | | (0.52) | | (0.07) | | (0.85) | | (0.13) | | | | | |
| -0.36 | -0.03 | -0.07 | | -0.10 | -0.10 | | | | -0.28 | 8 | -95.07 | 208.81 | 5.03 | 0.01 |
| (0.26) | (0.33) | (0.14) | | (0.31) | (0.07) | | | | (0.13) | | | | | |
| -0.32 | -0.03 | 0.01 | | | -0.17 | -0.24 | | | | 7 | -96.73 | 209.50 | 5.72 | 0.01 |
| (0.28) | (0.38) | (0.15) | | | (0.06) | (0.22) | | | | | | | | |
| -0.94 | 0.25 | 0.25 | 0.98 | | -0.15 | | | | | 7 | -96.88 | 209.80 | 6.02 | 0.01 |
| (0.52) | (0.31) | (0.32) | (0.96) | | (0.06) | | | | | | | | | |
| -0.62 | 0.25 | | 0.51 | -0.20 | -0.06 | | | -0.55 | -0.30 | 9 | -94.31 | 210.01 | 6.23 | 0.01 |
| (0.34) | (0.38) | | (0.44) | (0.31) | (0.07) | | | (0.51) | (0.13) | | | | | |
| -0.52 | 0.09 | | 0.20 | | -0.16 | | 0.22 | | | 7 | -97.12 | 210.28 | 6.49 | 0.01 |
| (0.44) | (0.80) | | (0.53) | | (0.06) | | (0.86) | | | | | | | |
| -0.58 | 0.27 | | 0.29 | -0.07 | -0.15 | | | | | 7 | -97.13 | 210.29 | 6.51 | 0.01 |
| (0.34) | (0.32) | | (0.44) | (0.32) | (0.06) | | | | | | | | | |
| -0.86 | | 0.28 | 0.88 | -0.10 | -0.16 | | | | | 7 | -97.14 | 210.32 | 6.54 | 0.01 |
| (0.53) | | (0.31) | (0.95) | (0.31) | (0.06) | | | | | | | | | |
| -0.46 | 0.23 | | | -0.07 | -0.17 | | | -0.22 | | 7 | -97.23 | 210.49 | 6.71 | 0.01 |
| (0.27) | (0.31) | | | (0.32) | (0.06) | | | (0.52) | | | | | | |
| -0.44 | 0.20 | -0.03 | | -0.05 | -0.16 | | | | | 7 | -97.29 | 210.62 | 6.84 | 0.01 |
| (0.27) | (0.32) | (0.15) | | (0.32) | (0.06) | | | | | | | | | |
| -0.37 | 0.15 | -0.10 | | -0.17 | -0.08 | | | -0.47 | -0.30 | 9 | -94.66 | 210.71 | 6.93 | 0.00 |
| (0.26) | (0.38) | (0.14) | | (0.31) | (0.07) | | | (0.51) | (0.13) | | | | | |
| **MEDIUM-TERM (PRIME AGE, FGMt, N = 90)** | | | | | | | | | | | | | | |
| **I** | **FGM** | **Qcoh** | **Pop(CH)** | **Sex(M)** | **Mass** | **FGM:Qcoh** | **FGM:Pop(CH)** | **FGM:Sex(M)** | **FGM:Mass** | **df** | **Log-lik** | **AICc** | **Delta** | **Weight** |
| -0.19 | | | | | -0.09 | | | | | 4 | -145.49 | 299.46 | 0.00 | 0.12 |
| (0.21) | | | | | (0.05) | | | | | | | | | |
| **-0.19** | | | | | | | | | | **3** | **-146.97** | **300.23** | **0.77** | **0.08** |
| **(0.21)** | | | | | | | | | | | | | | |
| -0.02 | | | -0.37 | | -0.12 | | | | | 5 | -144.96 | 300.64 | 1.19 | 0.07 |
| (0.26) | | | (0.34) | | (0.06) | | | | | | | | | |
| -0.19 | 0.14 | | | | -0.09 | | | | | 5 | -145.03 | 300.77 | 1.31 | 0.06 |
| (0.21) | (0.14) | | | | (0.05) | | | | | | | | | |
| -0.20 | | 0.07 | | | -0.11 | | | | | 5 | -145.25 | 301.22 | 1.76 | 0.05 |
| (0.21) | | (0.10) | | | (0.06) | | | | | | | | | |
| -0.24 | | | | 0.12 | -0.10 | | | | | 5 | -145.40 | 301.50 | 2.05 | 0.04 |
| (0.24) | | | | (0.28) | (0.06) | | | | | | | | | |

| | | | | | | | | | | | | | | |
|---|---|---|---|---|---|---|---|---|---|---|---|---|---|---|
| -0.19 | 0.13 | | | | | | | | | 4 | -146.58 | 301.63 | 2.17 | 0.04 |
| (0.21) | (0.15) | | | | | | | | | | | | | |
| -0.19 | | -0.04 | | | | | | | | 4 | -146.88 | 302.24 | 2.78 | 0.03 |
| (0.21) | | (0.09) | | | | | | | | | | | | |
| -0.16 | | | | -0.09 | | | | | | 4 | -146.91 | 302.28 | 2.83 | 0.03 |
| (0.23) | | | | (0.26) | | | | | | | | | | |
| -0.22 | | | 0.05 | | | | | | | 4 | -146.96 | 302.39 | 2.93 | 0.03 |
| (0.26) | | | (0.28) | | | | | | | | | | | |
| -0.18 | 0.11 | | | | -0.09 | | | | -0.04 | 6 | -144.72 | 302.46 | 3.00 | 0.03 |
| (0.22) | (0.15) | | | | (0.05) | | | | (0.05) | | | | | |
| -0.08 | | | -0.41 | 0.18 | -0.14 | | | | | 6 | -144.77 | 302.55 | 3.09 | 0.03 |
| (0.27) | | | (0.34) | (0.28) | (0.07) | | | | | | | | | |
| -0.07 | 0.09 | | -0.27 | | -0.11 | | | | | 6 | -144.80 | 302.62 | 3.16 | 0.02 |
| (0.27) | (0.16) | | (0.37) | | (0.06) | | | | | | | | | |
| 0.11 | | -0.09 | -0.63 | | -0.12 | | | | | 6 | -144.88 | 302.78 | 3.32 | 0.02 |
| (0.41) | | (0.21) | (0.69) | | (0.06) | | | | | | | | | |
| -0.20 | 0.12 | 0.04 | | | -0.10 | | | | | 6 | -144.97 | 302.94 | 3.49 | 0.02 |
| (0.21) | (0.16) | (0.11) | | | (0.06) | | | | | | | | | |
| -0.22 | 0.13 | | | 0.07 | -0.09 | | | | | 6 | -145.00 | 303.01 | 3.55 | 0.02 |
| (0.24) | (0.15) | | | (0.28) | (0.06) | | | | | | | | | |
| -0.27 | | 0.09 | | 0.17 | -0.13 | | | | | 6 | -145.08 | 303.17 | 3.71 | 0.02 |
| (0.24) | | (0.10) | | (0.28) | (0.07) | | | | | | | | | |
| -0.18 | 0.17 | -0.07 | | | | | | | | 5 | -146.28 | 303.28 | 3.82 | 0.02 |
| (0.22) | (0.15) | (0.09) | | | | | | | | | | | | |
| -0.13 | 0.15 | | | -0.15 | | | | | | 5 | -146.42 | 303.55 | 4.09 | 0.02 |
| (0.23) | (0.15) | | | (0.26) | | | | | | | | | | |
| -0.27 | 0.16 | | 0.18 | | | | | | | 5 | -146.43 | 303.57 | 4.11 | 0.02 |
| (0.27) | (0.16) | | (0.30) | | | | | | | | | | | |
| 0.01 | | -0.14 | -0.40 | | | | | | | 5 | -146.74 | 304.20 | 4.74 | 0.01 |
| (0.42) | | (0.21) | (0.70) | | | | | | | | | | | |
| -0.04 | 0.05 | | -0.30 | | -0.12 | | | | -0.05 | 7 | -144.44 | 304.25 | 4.79 | 0.01 |
| (0.28) | (0.16) | | (0.37) | | (0.06) | | | | (0.05) | | | | | |
| -0.15 | | -0.03 | | -0.08 | | | | | | 5 | -146.83 | 304.38 | 4.92 | 0.01 |
| (0.24) | | (0.09) | | (0.26) | | | | | | | | | | |
| -0.17 | | | 0.04 | -0.09 | | | | | | 5 | -146.90 | 304.51 | 5.05 | 0.01 |
| (0.28) | | | (0.29) | (0.26) | | | | | | | | | | |
| -0.20 | 0.22 | | | 0.09 | -0.09 | | | -0.27 | | 7 | -144.60 | 304.56 | 5.11 | 0.01 |
| (0.25) | (0.18) | | | (0.28) | (0.05) | | | (0.30) | | | | | | |
| -0.19 | 0.08 | 0.06 | | | -0.11 | | | | -0.05 | 7 | -144.60 | 304.57 | 5.11 | 0.01 |
| (0.21) | (0.16) | (0.11) | | | (0.06) | | | | (0.05) | | | | | |
| -0.22 | 0.09 | | | 0.10 | -0.10 | | | | -0.05 | 7 | -144.66 | 304.68 | 5.23 | 0.01 |
| (0.25) | (0.15) | | | (0.29) | (0.05) | | | | (0.05) | | | | | |
| -0.00 | -0.05 | | -0.32 | | -0.12 | | 0.19 | | | 7 | -144.67 | 304.71 | 5.25 | 0.01 |
| (0.30) | (0.32) | | (0.38) | | (0.06) | | (0.36) | | | | | | | |
| -0.10 | 0.07 | | -0.33 | 0.14 | -0.13 | | | | | 7 | -144.70 | 304.76 | 5.30 | 0.01 |
| (0.28) | (0.17) | | (0.39) | (0.30) | (0.07) | | | | | | | | | |

| | | | | | | | | | | | | | | |
|---|---|---|---|---|---|---|---|---|---|---|---|---|---|---|
| 0.04 | | -0.07 | -0.63 | 0.17 | -0.14 | | | | | 7 | -144.71 | 304.78 | 5.32 | 0.01 |
| (0.42) | | (0.21) | (0.69) | (0.29) | (0.07) | | | | | | | | | |
| 0.06 | 0.09 | -0.09 | -0.52 | | -0.11 | | | | | 7 | -144.72 | 304.81 | 5.36 | 0.01 |
| (0.41) | (0.16) | (0.21) | (0.70) | | (0.06) | | | | | | | | | |
| -0.12 | 0.24 | | | -0.13 | | | | -0.28 | | 6 | -146.02 | 305.05 | 5.59 | 0.01 |
| (0.24) | (0.18) | | | (0.26) | | | | (0.31) | | | | | | |
| -0.17 | 0.08 | 0.05 | | | -0.10 | -0.05 | | | | 7 | -144.86 | 305.09 | 5.64 | 0.01 |
| (0.22) | (0.17) | (0.11) | | | (0.06) | (0.11) | | | | | | | | |
| -0.24 | 0.10 | 0.05 | | 0.11 | -0.12 | | | | | 7 | -144.90 | 305.16 | 5.71 | 0.01 |
| (0.24) | (0.16) | (0.12) | | (0.30) | (0.07) | | | | | | | | | |
| -0.12 | 0.19 | -0.07 | | -0.14 | | | | | | 6 | -146.14 | 305.29 | 5.83 | 0.01 |
| (0.24) | (0.16) | (0.09) | | (0.26) | | | | | | | | | | |
| -0.15 | 0.14 | -0.06 | | | | -0.05 | | | | 6 | -146.19 | 305.39 | 5.93 | 0.01 |
| (0.23) | (0.17) | (0.09) | | | | (0.11) | | | | | | | | |
| -0.05 | 0.16 | -0.14 | -0.25 | | | | | | | 6 | -146.23 | 305.47 | 6.01 | 0.01 |
| (0.42) | (0.16) | (0.21) | (0.70) | | | | | | | | | | | |
| -0.21 | 0.18 | | 0.17 | -0.14 | | | | | | 6 | -146.28 | 305.57 | 6.11 | 0.01 |
| (0.29) | (0.16) | | (0.30) | (0.26) | | | | | | | | | | |
| -0.23 | 0.08 | | 0.16 | | | | 0.11 | | | 6 | -146.38 | 305.77 | 6.32 | 0.01 |
| (0.29) | (0.32) | | (0.31) | | | | (0.36) | | | | | | | |
| -0.08 | 0.01 | | -0.39 | 0.19 | -0.14 | | | | -0.05 | 8 | -144.24 | 306.26 | 6.80 | 0.00 |
| (0.28) | (0.18) | | (0.39) | (0.30) | (0.07) | | | | (0.06) | | | | | |
| 0.06 | | -0.15 | -0.42 | -0.09 | | | | | | 6 | -146.67 | 306.36 | 6.90 | 0.00 |
| (0.44) | | (0.21) | (0.70) | (0.26) | | | | | | | | | | |
| **MEDIUM-TERM (PRIME AGED, mean FGM, N = 71)** | | | | | | | | | | | | | | |
| **I** | **FGM** | **Qcoh** | **Pop(CH)** | **Sex(M)** | **Mass** | **FGM:Qcoh** | **FGM:Pop(CH)** | **FGM:Sex(M)** | **FGM:Mass** | **df** | **Log-lik** | **AICc** | **Delta** | **Weight** |
| **-0.28** | | | | | | | | | | **3** | **-120.25** | **246.85** | **0.00** | **0.12** |
| **(0.22)** | | | | | | | | | | | | | | |
| -0.28 | | | | | -0.08 | | | | | 4 | -119.38 | 247.37 | 0.52 | 0.09 |
| (0.23) | | | | | (0.06) | | | | | | | | | |
| -0.01 | | | -0.55 | | -0.13 | | | | | 5 | -118.52 | 247.96 | 1.11 | 0.07 |
| (0.29) | | | (0.39) | | (0.07) | | | | | | | | | |
| -0.30 | | 0.12 | | | -0.12 | | | | | 5 | -118.88 | 248.69 | 1.84 | 0.05 |
| (0.22) | | (0.12) | | | (0.07) | | | | | | | | | |
| -0.28 | -0.09 | | | | | | | | | 4 | -120.17 | 248.94 | 2.09 | 0.04 |
| (0.22) | (0.23) | | | | | | | | | | | | | |
| -0.22 | | | -0.13 | | | | | | | 4 | -120.18 | 248.97 | 2.12 | 0.04 |
| (0.28) | | | (0.34) | | | | | | | | | | | |
| -0.24 | | | | -0.11 | | | | | | 4 | -120.19 | 248.98 | 2.13 | 0.04 |
| (0.26) | | | | (0.31) | | | | | | | | | | |
| -0.29 | | 0.01 | | | | | | | | 4 | -120.24 | 249.09 | 2.24 | 0.04 |
| (0.22) | | (0.10) | | | | | | | | | | | | |
| 0.14 | -0.30 | | -0.82 | | -0.15 | | | | | 6 | -117.89 | 249.10 | 2.25 | 0.04 |
| (0.31) | (0.26) | | (0.45) | | (0.07) | | | | | | | | | |
| -0.34 | | | | 0.13 | -0.09 | | | | | 5 | -119.31 | 249.54 | 2.69 | 0.03 |
| (0.27) | | | | (0.35) | (0.07) | | | | | | | | | |

| | | | | | | | | | | | | | | |
|---|---|---|---|---|---|---|---|---|---|---|---|---|---|---|
| -0.28 | -0.07 | | | | -0.08 | | | | | 5 | -119.33 | 249.59 | 2.74 | 0.03 |
| (0.23) | (0.23) | | | | (0.06) | | | | | | | | | |
| -0.08 | | | -0.60 | 0.23 | -0.16 | | | | | 6 | -118.32 | 249.95 | 3.10 | 0.03 |
| (0.31) | | | (0.40) | (0.36) | (0.08) | | | | | | | | | |
| 0.12 | | -0.08 | -0.78 | | -0.13 | | | | | 6 | -118.47 | 250.25 | 3.40 | 0.02 |
| (0.49) | | (0.24) | (0.80) | | (0.07) | | | | | | | | | |
| -0.31 | -0.23 | 0.18 | | | -0.13 | | | | | 6 | -118.49 | 250.29 | 3.44 | 0.02 |
| (0.21) | (0.26) | (0.14) | | | (0.07) | | | | | | | | | |
| 0.06 | -0.37 | | -0.97 | 0.36 | -0.20 | | | | | 7 | -117.40 | 250.58 | 3.73 | 0.02 |
| (0.32) | (0.27) | | (0.46) | (0.37) | (0.09) | | | | | | | | | |
| -0.40 | | 0.14 | | 0.21 | -0.15 | | | | | 6 | -118.71 | 250.72 | 3.88 | 0.02 |
| (0.27) | | (0.12) | | (0.36) | (0.08) | | | | | | | | | |
| -0.16 | -0.17 | | -0.25 | | | | | | | 5 | -119.98 | 250.88 | 4.04 | 0.02 |
| (0.29) | (0.26) | | (0.37) | | | | | | | | | | | |
| 0.05 | | -0.16 | -0.63 | | | | | | | 5 | -120.00 | 250.92 | 4.07 | 0.02 |
| (0.50) | | (0.25) | (0.81) | | | | | | | | | | | |
| -0.16 | | | -0.15 | -0.12 | | | | | | 5 | -120.10 | 251.13 | 4.28 | 0.01 |
| (0.32) | | | (0.34) | (0.31) | | | | | | | | | | |
| -0.25 | -0.08 | | | -0.09 | | | | | | 5 | -120.13 | 251.18 | 4.33 | 0.01 |
| (0.26) | (0.24) | | | (0.31) | | | | | | | | | | |
| -0.29 | -0.13 | 0.03 | | | | | | | | 5 | -120.13 | 251.18 | 4.33 | 0.01 |
| (0.22) | (0.26) | (0.11) | | | | | | | | | | | | |
| -0.24 | | 0.01 | | -0.11 | | | | | | 5 | -120.18 | 251.28 | 4.43 | 0.01 |
| (0.26) | | (0.10) | | (0.31) | | | | | | | | | | |
| 0.17 | -0.32 | | -0.86 | | -0.16 | | | | -0.03 | 7 | -117.84 | 251.45 | 4.60 | 0.01 |
| (0.32) | (0.27) | | (0.46) | | (0.07) | | | | (0.08) | | | | | |
| 0.08 | -0.17 | | -0.77 | | -0.15 | | -0.16 | | | 7 | -117.86 | 251.50 | 4.65 | 0.01 |
| (0.38) | (0.59) | | (0.48) | | (0.07) | | (0.65) | | | | | | | |
| 0.22 | -0.29 | -0.05 | -0.97 | | -0.15 | | | | | 7 | -117.87 | 251.52 | 4.67 | 0.01 |
| (0.49) | (0.26) | (0.24) | (0.80) | | (0.07) | | | | | | | | | |
| -0.34 | -0.08 | | | 0.15 | -0.09 | | | | | 6 | -119.25 | 251.80 | 4.95 | 0.01 |
| (0.28) | (0.23) | | | (0.35) | (0.07) | | | | | | | | | |
| -0.28 | -0.07 | | | | -0.08 | | | | 0.01 | 6 | -119.33 | 251.98 | 5.13 | 0.01 |
| (0.23) | (0.24) | | | | (0.06) | | | | (0.08) | | | | | |
| -0.46 | -0.30 | 0.22 | | 0.32 | -0.18 | | | | | 7 | -118.11 | 251.99 | 5.14 | 0.01 |
| (0.27) | (0.27) | (0.14) | | (0.37) | (0.09) | | | | | | | | | |
| -0.38 | -0.17 | 0.16 | | | -0.13 | 0.11 | | | | 7 | -118.27 | 252.32 | 5.47 | 0.01 |
| (0.23) | (0.28) | (0.14) | | | (0.07) | (0.17) | | | | | | | | |
| 0.03 | | -0.06 | -0.79 | 0.22 | -0.15 | | | | | 7 | -118.28 | 252.35 | 5.50 | 0.01 |
| (0.51) | | (0.25) | (0.79) | (0.36) | (0.08) | | | | | | | | | |
| -0.31 | -0.26 | 0.19 | | | -0.14 | | | | -0.02 | 7 | -118.45 | 252.67 | 5.82 | 0.01 |
| (0.22) | (0.27) | (0.14) | | | (0.08) | | | | (0.09) | | | | | |
| 0.10 | -0.43 | | -1.06 | 0.41 | -0.21 | | | | -0.05 | 8 | -117.24 | 252.80 | 5.95 | 0.01 |
| (0.33) | (0.29) | | (0.49) | (0.37) | (0.09) | | | | (0.09) | | | | | |
| 0.10 | -0.16 | -0.15 | -0.72 | | | | | | | 6 | -119.81 | 252.93 | 6.08 | 0.01 |
| (0.50) | (0.26) | (0.25) | (0.81) | | | | | | | | | | | |

| | | | | | | | | | | | | | | |
|---|---|---|---|---|---|---|---|---|---|---|---|---|---|---|
| 0.05 | -0.43 | | -0.97 | 0.36 | -0.20 | | | 0.17 | | 8 | -117.34 | 253.00 | 6.15 | 0.01 |
| (0.32) | (0.31) | | (0.46) | (0.37) | (0.09) | | | (0.48) | | | | | | |
| 0.01 | -0.25 | | -0.92 | 0.36 | -0.20 | | -0.15 | | | 8 | -117.37 | 253.07 | 6.22 | 0.01 |
| (0.38) | (0.59) | | (0.50) | (0.37) | (0.09) | | (0.65) | | | | | | | |
| -0.25 | 0.05 | | -0.17 | | | | -0.27 | | | 6 | -119.90 | 253.10 | 6.26 | 0.01 |
| (0.36) | (0.60) | | (0.41) | | | | (0.66) | | | | | | | |
| 0.10 | -0.37 | -0.02 | -1.04 | 0.36 | -0.20 | | | | | 8 | -117.40 | 253.12 | 6.27 | 0.01 |
| (0.51) | (0.27) | (0.25) | (0.80) | (0.37) | (0.09) | | | | | | | | | |
| -0.36 | -0.06 | 0.01 | | | | 0.11 | | | | 6 | -119.91 | 253.13 | 6.28 | 0.01 |
| (0.24) | (0.28) | (0.12) | | | | (0.17) | | | | | | | | |
| 0.11 | | -0.16 | -0.64 | -0.12 | | | | | | 6 | -119.92 | 253.15 | 6.30 | 0.01 |
| (0.52) | | (0.25) | (0.81) | (0.31) | | | | | | | | | | |
| -0.11 | -0.16 | | -0.26 | -0.11 | | | | | | 6 | -119.92 | 253.16 | 6.31 | 0.01 |
| (0.33) | (0.26) | | (0.38) | (0.31) | | | | | | | | | | |
| -0.25 | -0.12 | 0.04 | | -0.10 | | | | | | 6 | -120.08 | 253.47 | 6.62 | 0.00 |
| (0.26) | (0.26) | (0.11) | | (0.31) | | | | | | | | | | |
| -0.25 | -0.11 | | | -0.09 | | | | 0.10 | | 6 | -120.11 | 253.53 | 6.68 | 0.00 |
| (0.26) | (0.28) | | | (0.31) | | | | (0.49) | | | | | | |
| 0.13 | -0.24 | -0.05 | -0.89 | | -0.15 | 0.08 | | | | 8 | -117.77 | 253.85 | 7.00 | 0.00 |
| (0.53) | (0.28) | (0.24) | (0.81) | | (0.07) | (0.17) | | | | | | | | |

Table S4. Linear mixed effects model (LMM) selection table for the long-term relationship between adult body mass and faecal glucocorticoid metabolites ('FGM') measured as juvenile. We accounted for cohort quality ('Qcoh'), population (reference level is Chizé: CH, and is compared to Trois-Fontaines: TF), and sex (reference level is male: M, and is compared to females: F). Values give the parameter coefficient and values between brackets in grey rows are the standard-errors. 'I' is the Intercept, 'df' is the number of parameters, 'Log-lik' is the log-likelihood, 'delta' is the difference of AICc between the candidate model and the model having the lowest AICc, and 'weight' the AIC weight of each model. Retained model is in bold. Only models up to delta AICc = 7 are displayed.

| **LONG-TERM (N = 345 obs., n = 159 ind.)** | | | | | | | | | | | | |
|---|---|---|---|---|---|---|---|---|---|---|---|---|
| **I** | **FGM** | **Qcoh** | **Pop(CH)** | **Sex(M)** | **FGM:Qcoh** | **FGM:Pop(CH)** | **FGM:Sex(M)** | **df** | **Log-lik** | **AICc** | **Delta** | **Weight** |
| -0.04 | | 0.40 | | | | | | 5 | -652.34 | 1314.85 | 0.00 | 0.20 |
| (0.28) | | (0.20) | | | | | | | | | | |
| -0.20 | | 0.41 | 0.33 | | | | | 6 | -651.82 | 1315.90 | 1.05 | 0.12 |
| (0.32) | | (0.20) | (0.32) | | | | | | | | | |
| 0.04 | | 0.39 | | -0.17 | | | | 6 | -652.20 | 1316.65 | 1.80 | 0.08 |
| (0.32) | | (0.20) | | (0.32) | | | | | | | | |
| **-0.06** | | | | | | | | **4** | **-654.27** | **1316.66** | **1.81** | **0.08** |
| **(0.29)** | | | | | | | | | | | | |
| -0.04 | -0.04 | 0.40 | | | | | | 6 | -652.32 | 1316.88 | 2.03 | 0.07 |
| (0.28) | (0.19) | (0.20) | | | | | | | | | | |
| -0.10 | | 0.40 | 0.35 | -0.20 | | | | 7 | -651.64 | 1317.61 | 2.76 | 0.05 |
| (0.35) | | (0.20) | (0.33) | (0.32) | | | | | | | | |
| -0.21 | | | 0.32 | | | | | 5 | -653.80 | 1317.77 | 2.92 | 0.05 |
| (0.33) | | | (0.33) | | | | | | | | | |
| -0.20 | 0.02 | 0.41 | 0.34 | | | | | 7 | -651.82 | 1317.97 | 3.12 | 0.04 |

| | | | | | | | | | | | | |
|---|---|---|---|---|---|---|---|---|---|---|---|---|
| (0.33) | (0.20) | (0.20) | (0.34) | | | | | | | | | |
| 0.05 | | | | -0.23 | | | | 5 | -654.03 | 1318.23 | 3.37 | 0.04 |
| (0.33) | | | | (0.32) | | | | | | | | |
| -0.06 | -0.04 | | | | | | | 5 | -654.25 | 1318.68 | 3.82 | 0.03 |
| (0.29) | (0.19) | | | | | | | | | | | |
| 0.05 | -0.04 | 0.39 | | -0.17 | | | | 7 | -652.18 | 1318.70 | 3.85 | 0.03 |
| (0.32) | (0.19) | (0.20) | | (0.32) | | | | | | | | |
| -0.04 | -0.04 | 0.41 | | | -0.09 | | | 7 | -652.26 | 1318.85 | 4.00 | 0.03 |
| (0.28) | (0.19) | (0.20) | | | (0.26) | | | | | | | |
| -0.09 | | | 0.34 | -0.26 | | | | 6 | -653.49 | 1319.22 | 4.37 | 0.02 |
| (0.36) | | | (0.33) | (0.33) | | | | | | | | |
| -0.11 | 0.03 | 0.40 | 0.36 | -0.20 | | | | 8 | -651.63 | 1319.69 | 4.84 | 0.02 |
| (0.36) | (0.20) | (0.20) | (0.34) | (0.32) | | | | | | | | |
| -0.22 | 0.02 | | 0.33 | | | | | 6 | -653.80 | 1319.84 | 4.99 | 0.02 |
| (0.33) | (0.20) | | (0.34) | | | | | | | | | |
| -0.20 | 0.02 | 0.42 | 0.34 | | -0.10 | | | 8 | -651.75 | 1319.94 | 5.08 | 0.02 |
| (0.33) | (0.20) | (0.20) | (0.34) | | (0.26) | | | | | | | |
| -0.19 | -0.03 | 0.41 | 0.33 | | | 0.07 | | 8 | -651.81 | 1320.04 | 5.19 | 0.01 |
| (0.34) | (0.35) | (0.20) | (0.35) | | | (0.42) | | | | | | |
| 0.04 | 0.11 | 0.40 | | -0.14 | | | -0.29 | 8 | -651.89 | 1320.21 | 5.36 | 0.01 |
| (0.32) | (0.27) | (0.20) | | (0.32) | | | (0.37) | | | | | |
| 0.06 | -0.04 | | | -0.23 | | | | 6 | -654.01 | 1320.26 | 5.41 | 0.01 |
| (0.33) | (0.19) | | | (0.33) | | | | | | | | |
| 0.04 | -0.04 | 0.40 | | -0.16 | -0.08 | | | 8 | -652.14 | 1320.71 | 5.85 | 0.01 |
| (0.32) | (0.19) | (0.21) | | (0.32) | (0.26) | | | | | | | |
| -0.13 | 0.19 | 0.41 | 0.38 | -0.17 | | | -0.31 | 9 | -651.28 | 1321.10 | 6.24 | 0.01 |
| (0.36) | (0.28) | (0.20) | (0.34) | (0.32) | | | (0.37) | | | | | |
| -0.10 | 0.03 | | 0.36 | -0.26 | | | | 7 | -653.48 | 1321.29 | 6.44 | 0.01 |
| (0.37) | (0.20) | | (0.35) | (0.33) | | | | | | | | |
| -0.12 | 0.02 | 0.40 | 0.36 | -0.19 | -0.08 | | | 9 | -651.58 | 1321.70 | 6.84 | 0.01 |
| (0.36) | (0.20) | (0.21) | (0.34) | (0.32) | (0.26) | | | | | | | |
| -0.09 | -0.03 | 0.39 | 0.34 | -0.20 | | 0.08 | | 9 | -651.61 | 1321.76 | 6.91 | 0.01 |
| (0.37) | (0.35) | (0.20) | (0.35) | (0.32) | | (0.42) | | | | | | |
